# Hawkes process modelling for chemical reaction networks in a random environment

**DOI:** 10.1101/2023.08.25.554803

**Authors:** Mark Sinzger-D’Angelo, Heinz Koeppl

## Abstract

Cellular processes are open systems, situated in a heterogeneous context, rather than operating in isolation. Chemical reaction networks (CRNs) whose reaction rates are modelled as external stochastic processes account for the heterogeneous environment when describing the embedded process. A marginal description of the embedded process is of interest for (i) fast simulations that bypass the co-simulation of the environment, (ii) obtaining new process equations from which moment equations can be derived, (iii) the computation of information-theoretic quantities, and (iv) state estimation. It is known since Snyder’s and related works that marginalization over a stochastic intensity turns point processes into self-exciting ones. While the Snyder filter specifies the exact history-dependent propensities in the framework of CRNs in Markov environment, it was recently suggested to use approximate filters for the marginal description. By regarding the chemical reactions as events, we establish a link between CRNs in a linear random environment and Hawkes processes, a class of self-exciting counting processes widely used in event analysis. The Hawkes approximation can be obtained via moment closure scheme or as the optimal linear approximation under the quadratic criterion. We show the equivalence of both approaches. Furthermore, we use martingale techniques to provide results on the agreement of the Hawkes process and the exact marginal process in their second order statistics, i.e., covariance, auto/cross-correlation. We introduce an approximate marginal simulation algorithm and illustrate it in case studies.

**AMS subject classifications:** 37M05, 60G35, 60G55, 60J28, 60K37, 62M15

## 1 Introduction

Stochastic chemical reaction networks (CRNs) model the synthesis, conversion and decay of molecules under the assumption of spatial homogeneity and dominant effect of low copy numbers [39]. Often the processes that are modelled are not closed, but are embedded in the cellular context, which we call random environment [57]. The concept of the random environment is abstract enough to include cell-to-cell heterogeneity, such as a varying polymerase copy number, as well as fluctuating processes within the cell, such as conformation changes of mRNA. One way to model the embedding into a random environment is by making the propensity functions depend on an external stochastic process. In other words, the reaction rate constants are replaced by random (time-varying) quantities. An embedding into a random environment naturally arises when one part of a larger CRN is labelled as the environment and the remaining part is the process of interest. This situation is typically encountered in the experimental context when a network is only partially observed. Diffusion processes that remain positive, such as the Cox-Ingersoll-Ross process [12] or a positive function of an Ornstein-Uhlenbeck process, are other ways to model fluctuating concentrations as environment components.

It is one modelling goal to find a closed, i.e., self-contained description for a given CRN in a random environment that behaves in its statistical properties as if it was still embedded. We call it the marginal description. To achieve this goal, previous works established the intimate link to stochastic filtering [56, 9]. The term stochastic filtering is a synonym for dynamic Bayesian estimation of the random environment components from the observed part of the network [7]. The posterior mean (or filter mean) of the random propensity conditional on the full history of observed reactions is the central quantity that is needed for the marginal description. However, computing it is often impossible. For this reason, we look for approximations of the filter mean. Approximate filters are typically obtained from projections, variational methods, moments closures and assumed density approaches [57, 58, 9].

There are various purposes for studying the filter mean and approximate filters, which we summarize in this section. First, we have already motivated the study for model reduction purposes. Along this line, Zechner, Bronstein and Koeppl introduced the marginal process framework [56, 9] as a technical tool to reduce Markov jump models. For the reduced models, the Markov property on an augmented state space can be used to derive generalized (auxiliary-variable) [35, 50] or reduced [41] master equations. Second, the work by Zechner and Koeppl [57] proposed the marginal simulation algorithm that bypasses the co-simulation of environment components, emphasising its potential for efficient simulation and variance reduction [18] as known from Rao-Blackwellized estimators [46, 6].

Beyond the marginal description, the filter mean naturally appears in expressions for information-theoretic quantities. Shamai and Lapidoth used the optimal linear causal estimate to obtain upper bounds on the capacity for the spectrally constrained Poisson channel [49]. Duso, Moor and Zechner used the Gamma filter [19, 40] to estimate the mutual information between two species in a CRN. Furthermore, it has been known since the work on counting processes by Brémaud [7, §VI.6] that the relative entropy between two counting processes is a difference functional of the filter means (‘detection formula’). Atar and Weissman used it to derive the error of mismatched estimation of the mutual information, when using an approximate filter [3]. We presented a method to compute the relative entropy between counting processes and the mutual information of the Poisson by using piecewise-deterministic Markov processes [50].

Finally, to make this list complete, we emphasize that the original purpose for the study of the filter mean was state estimation under partial observability [52, 7]. In the literature on CRNs, this approach has been recently rediscovered. In their works, Rathinam et al. and Fang et al. introduce particle filters to solve the filtering problem for latent state estimation [47, 21]. State estimation for CRNs via smoothing, i.e., under the (partial) knowledge of the trajectory until the final time point, is addressed in [31]. The causal, anti-causal and non-causal estimator of the intensity, corresponding to forward filtering, backward filtering and smoothing are usually closely related, as in [27] for the Poisson process. Along with state estimation, one of the earliest application was in optimal control [7, §VII] sparked by the initial success of Kalman [33]. A more recent application designed genetic circuits using approximate filters [58].

After having clarified the various purposes of filtering theory for CRNs, we review the progress on approximate filters. The construction of approximate filters has been guided by the principle of moment closure and projection onto a tractable process class. Zechner used assumed density filtering with the Poisson and the Gamma assumption on the filtering distribution [58]. Eden and Brown as well as Harel and Opper used a Gaussian assumption [20, 29] for the state estimation from neural spike trains. Bronstein derived the method of entropic matching, which projects the filtering distribution along the Kullback-Leibler divergence time-point wise [9]. It was demonstrated for the product-Bernoulli and the product-Poisson ansatz. Alternatively, the minimization of the Kullback-Leibler divergence for the path likelihoods was recently proposed as a guiding principle for the model reduction [41]. This principle had been successfully applied for variational inference before [55].

In this paper, we pick up an idea of optimal linear filtering that Snyder [53] originally used for state estimation and recycle it for approximate marginal simulation. We establish a link with the self-exiting Hawkes counting process model used in a variety of domains, such as meteorology, finance and neuroscience. The Hawkes counting process model [30] is very well studied. The path likelihood for parameter estimation is known [44], simulation schemes are available [42] and recently Gao and Zhu [24] derived functional limit theorems with application to queues.

For the case in which only zeroth-order reactions are modulated, we derive that the approximation captures the proper second-order moment structure and show the equivalence with a moment-closure procedure. We derive the power spectral density for linear CRNs whose zeroth-order reactions are modulated by a correlated stationary environment. We contribute an approximate marginal simulation algorithm that is very efficient for independent environment components. We demonstrate our approximation method in three case studies. In these we highlight its strengths and limitations when it comes to bimodal distributions, the qualitative capturing of different timescales and the phenomenon of excursion due to periods of low values in the decay rate.

### 1.1 Counting process description of CRNs and the notion of sigma-algebras

CRNs formalize the stochastic dynamics of molecules of *n* types as Markov jump processes in the state space ℕ^*n*^ of species copy numbers on some probability space (Ω, *ℱ*, ℙ). The state *X*(*t*) = [*X*_1_(*t*), …, *X*_*n*_(*t*)]^*T*^ contains the number of molecules of type X_1_, …, X_*n*_ at time *t*. Transitions of the Markov jump process are dictated by *M* reactions

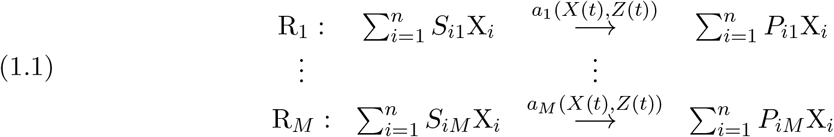

where *S*_*ij*_, *i* = 1, …, *n, j* = 1, …, *M* are the substrate coefficients and *P*_*ij*_, *i* = 1, …, *n, j* = 1, …, *M* the product coefficients. The superscripts *a*_*j*_(*X*(*t*), *Z*(*t*)) are the propensity functions. We assume that they depend on the current state *X*(*t*) and on an external stochastic process *Z*(*t*). By zeroth-order modulation, we denote a form *a*_*j*_(*x, z*) = *a*^*T*^ *z* + *b, a* ∈ ℝ^*l*^, *b* ∈ ℝ and by first-order modulation, we denote a form *a*_*j*_(*x, z*) = *x*^*T*^ *Az* + *x*^*T*^ *b, A* ∈ ℝ^*n×l*^, *b* ∈ ℝ^*n*^. We call *N* = *P − S* the stoichiometric matrix and its columns *ν*_1_, …, *ν*_*M*_ the change vectors. Transitions of the Markov jump process are possible from *x* to *x* + *ν*_*j*_ and occur at a rate given by the propensity function. We make this rigorous in the following counting process description. First, we introduce the multivariate counting process *Y* (*t*) = [*Y*_1_(*t*), …, *Y*_*M*_ (*t*)]^*T*^, where the component *Y*_*j*_(*t*) counts how often the Reaction R_*j*_ has occurred until time *t*, so naturally throughout *Y* (0) = 0. Then the current state *X*(*t*) can be computed from the initial state and *Y* (*t*) as

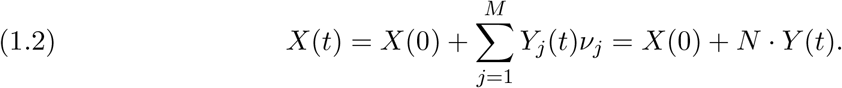

We use the terms point process and counting process equivalently. A filtration is an increasing family of sigma-algebras, indexed by time, that, intuitively speaking, captures our knowledge at time *t*. Define the filtrations

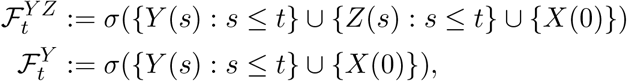

where *σ*(*·*) denotes the smallest sigma-algebra with respect to which all random variables in the argument are measurable. Then we can make the reaction system in Eq. (1.1) rigorous, by specifying that *Y* (*t*) has 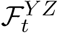 -intensity *λ*(*t*) = [*a*_1_(*X*(*t−*), *Z*(*t−*)), …, *a*_*M*_ (*X*(*t−*), *Z*(*t−*))]^*T*^. By *t−* we denote the left-sided limit, which makes *λ*(*t*) a predictable intensity. Later, we consider different probability measures, which we indicate by 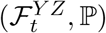 -intensity. For an introduction on intensities and their dependency on the filtration and the measure, we refer the reader to [7, §II]. Note, that, by Eq. (1.2), *a*_*j*_(*X*(*t−*), *Z*(*t−*)) is indeed 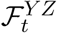 -measurable. In short, we write

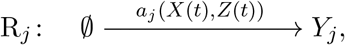

abusing the notation of birth processes for the reaction counters *Y*_1_, …, *Y*_*M*_. By Brémaud’s theorem on the change of filtration [7, §II, theorem T14], we obtain that the predictable 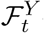 -intensity is 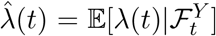, provided that a left-continuous version of the conditional expectation exists. This is always the case in our study, as we will see. We then write

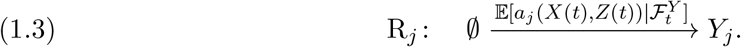

There are a couple of reasons why to study the intensity for the changed filtration. In the introduction, we highlighted (i) fast simulation, (ii) new process equations that give rise to new moment equations, (iii) the computation of information measures, and (iv) state estimation. Focusing on (i), we pick up the following idea of [57]. We use evolution equations that were obtained for 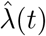, by stochastic filtering (see paragraph 1.5 below), to inform a stochastic simulation algorithm for *Y* (*t*) that handles time-dependent intensities. In this way, the equations, which were originally derived to estimate the environment state *Z*(*t*), are repurposed for marginal simulation of *Y* (*t*). Since, the filtering equations are impractical to work with, we use approximations thereof. We take Eq. (1.3) as the starting point for approximating (*Y* (*t*) ℙ) by (*Y* (*t*), ℚ) via an 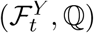 -intensity in Eq. (1.7)-(1.9) below. In order to see a prototypical evolution equation for 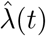, we first review a well-studied counting process.

### 1.2 Hawkes processes

In general, the 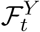 -intensity is some measurable function of the trajectory *Y*_[0,*t*]_ := {*Y* (*t*)}_0*≤s≤t*_ and *X*(0), by the very meaning of 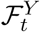 -measurable. In order to convey the intuition of how an 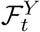 -intensity can be specified in practice, we consider the Hawkes counting process model in one dimension with exponential kernel. Its positive parameters are the base level *μ*, the jump height *β* and the decay *α*. In differential form, the 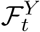 -intensity 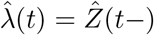 reads

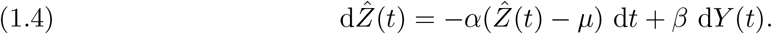

When a jump of *Y* occurs at time *t*, then d*Y* (*t*) = 1 and hence 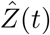 increases by *β*. In periods of no jumps, the intensity evolves according to the ODE specified in the first part of the evolution equation. In integral form it reads

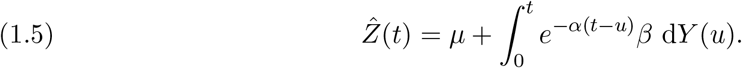

Under the assumption *α*^*−*1^*β <* 1, there exists a stationary version of *Y* (*t*) [8]. According to [13, §7.2, Example 7.2 (b)], the process can be defined on the entire real axis, replacing the lower integral bound by −*∞*.

In multiple dimensions, the Hawkes process with exponential kernel is more delicate. Hawkes [30] introduced the mutually exciting model for which the 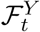 -intensity 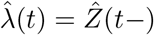 reads in coordinate integral form

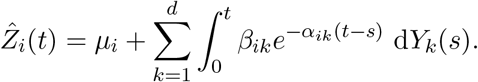

When a jump of *Y*_*k*_ occurs at time *t*, then d*Y*_*k*_(*t*) = 1 and hence 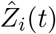 increases by *β*_*ik*_. In particular, a jump of a component can influence the intensity of another component, which is usually referred to as mutually exciting. It is different from reading Eq. (1.4) or Eq. (1.5) as matrix equations. Nevertheless, we call the matrix interpretation also Hawkes model, while being aware that this is not the classical exponential Hawkes model in multiple dimensions. Its coordinate differential form reads

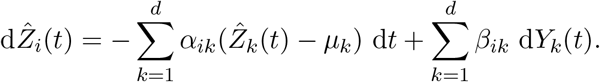

It generally requires taking the positive part of the intensity components 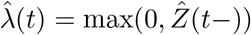, except in special cases where positivity can be guaranteed. This makes *Y* (*t*) a non-linear Hawkes model. The condition for the existence of a stationary version translates into *α* being invertible and *α*^*−*1^*β* having a spectral radius strictly less than 1.

### 1.3 Martingale calculus for counting processes

Associated with a (multivariate) counting process *Y* (*t*) ∈ ℕ^*d*^ and its *ℱ*_*t*_-intensity *λ*(*t*) for some filtration *ℱ*_*t*_ that contains *σ*(*Y* (*s*) : *s ≤ t*), there is the zero-mean *ℱ*_*t*_-martingale

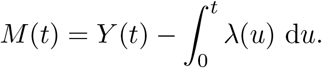

In fact, the property that the right-hand side is an *ℱ*_*t*_-martingale can be taken as the definition of the *ℱ*_*t*_-intensity for non-explosive *Y* (*t*). With this definition, it becomes apparent that the intensity depends on the filtration of the martingale. The increment can be written as d*M* (*t*) = d*Y* (*t*) *− λ*(*t*) d*t*. One benefit of martingale theory is that it provides a rich secondorder calculus. We name the identities that we will use, see for instance [1, proposition II.4.1]. For predictable processes 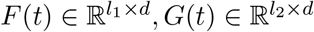, we have

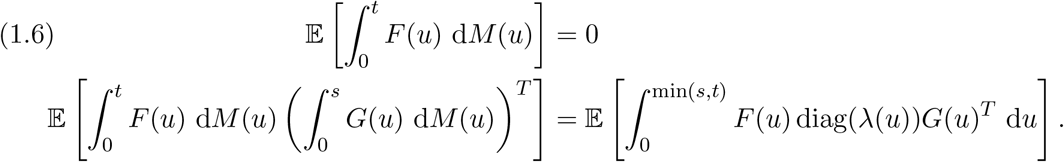

### 1.4 Standard conditions

In most of the remaining paper, we make simplifying assumptions on the external process *Z*(*t*) ∈ ℝ^*l*^ and the functional form *a*(*X*(*t*), *Z*(*t*)) of the 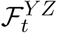 -intensity *λ*(*t*). We refer to these as the standard conditions. For the process *Z*(*t*), we assume

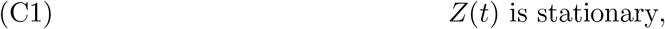

there exist *A*, ∑ ∈ ℝ^*l×l*^ such that all eigenvalues of *A* have positive real part, Σ is symmetric positive semi-definite, *A*Σ + Σ*A*^*T*^ is positive semi-definite and

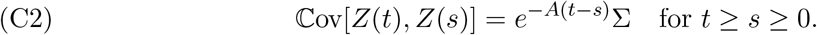

For the form of *λ*(*t*) = *a*(*X*(*t*), *Z*(*t*)) we assume

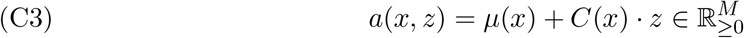

for some *μ*(*x*) ∈ ℝ^*M*^, *C*(*x*) ∈ ℝ^*M ×l*^. These simplifying assumptions include all linear reaction networks *Z*(*t*) with stable equilibrium, i.e., *Z*(*t*) is a stationary CRNs with only zeroth- and first-order reactions. For instance, this class covers the random telegraph model, compartmental models with only monomolecular reactions as well as cascades of birth-death processes. Gardiner’s regression theorem [25, §3.7.4b, Eq. (3.7.62), p.65] guarantees the matrix exponential form of the time correlation matrix. For processes with continuous state space, the Cox-Ingersoll-Ross process [12] satisfies these conditions. All the examples are Markov processes. For later use in the results section, we consider another condition

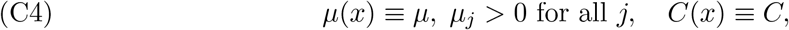

i.e., the propensities do not depend on the state *X*(*t*). This is the case, when only zeroth-order reactions are modulated by the environment and *Z*(*t*) is of mean zero.

### 1.5 Snyder’s exact filter

We now turn towards the study of the 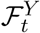 -intensity 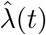, Eq. (1.3). Finding the posterior mean is a problem of Bayesian estimation, and filtering theory is concerned with its solution in the dynamical setting. We assume that (C1) holds and additionally that *Z*(*t*) is a Markov process. Then the Snyder filter specifies the evolution of 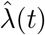. It can be formulated in terms of the posterior probabilities 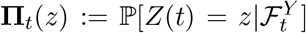 for *Z*(*t*) on a discrete state space and **Π**_*t*_(*z*) the respective posterior probability density on a continuous state space. Let the prior probability *p*_*t*_(*z*) evolve according to the operator *A*, i.e.,

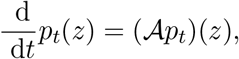

which can be the chemical master equation, if *Z*(*t*) is itself a CRN, or the Fokker-Planck equation, if *Z*(*t*) is a diffusion process given by a stochastic differential equation. Then, extending [53, Eq. (7.138), p.392 & Eq. (7.151), p.396], the posterior probabilities evolve as

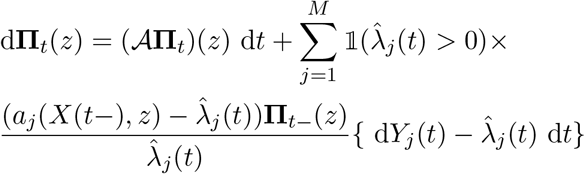

and the posterior moment is 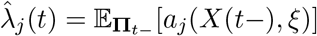, where we mean that *ξ* is distributed according to **Π**_*t−*_. The evolution equation is initialized in the stationary (prior) distribution of *Z*(*t*). The term after the prior dynamics is called the innovation term. It is the 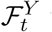 -martingale increment scaled by a predictable factor, the innovation gain. This structure ensures that the expectation of the posterior probability 𝔼[**Π**_*t*_] is constantly the stationary (prior) distribution of *Z*(*t*).

Assume additionally, that (C3) holds, and denote by *F* and *G* the prior dynamics of the mean and the covariance of *Z*(*t*), for which we assume they are functions of the first, second and third prior moments

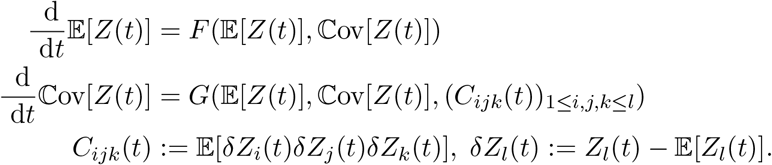

Then the evolution of the posterior mean 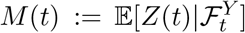 can be expressed with the auxiliary processes, which are the second- and third-order posterior moments

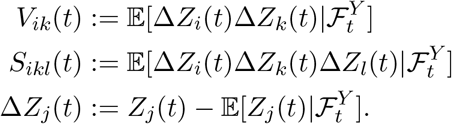

It reads

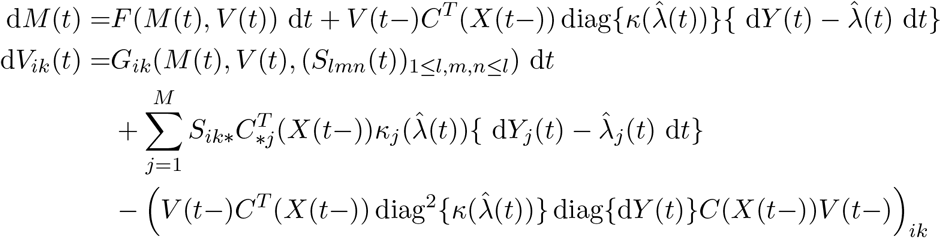

where 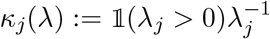 and 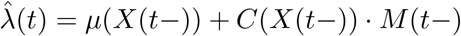

For more details on filtering of point processes, see [7, §IV] and for the more explicit setting of CRNs, we refer the reader to [57] and [9].

In general, the posterior moment equations are not closed and the evolution of the posterior probabilities can become prohibitively high-dimensional. Approximations to the evolution of the posterior mean are obtained by projection onto a tractable subclass of stochastic processes, which is achieved by minimizing a distance measure. The subclass can be defined by a sufficient statistics of low dimension or by a variational function. As an alternative ad-hoc approach to projection, posterior moment closures are employed. Assuming that the posterior distribution belongs to a parametric family, moment relations can be used to express higher order posterior moments by lower order ones. This closes the hierarchy of posterior moment equations.

### 1.6 The optimal linear filter

In the previous paragraph, we have discussed the problem of unclosed posterior moment equations. Here, we consider a particular projection method to obtain an approximate filter, the optimal linear filter 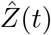. It was originally introduced for state estimation of the external signal *Z*(*t*) that modulates a doubly stochastic Poisson process [53, Eq. (7.238), p.429]. By optimal it is meant that it minimizes the quadratic criterion, i.e., it is the solution

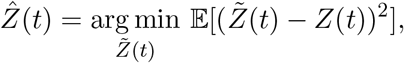

among all estimators

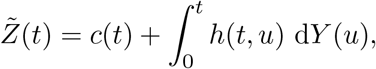

i.e., affine-linear in the trajectory *Y*_[0,*t*]_. The doubly stochastic Poisson process precisely corresponds to zeroth-order modulation. In case of higher order modulation we generalize the optimal linear estimator which results in *c*(*t*) and *h*(*t, u*) with a dependency on the history *X*_[0,*t*)_. Assuming the standard conditions (C1)-(C3), the optimal *c*(*t*) and *h*(*t, u*) can be conveniently formulated in differential vector form with an auxiliary Riccati equation [53, Eq. (7.238), p.429]. By shifting *Z*(*t*) to *Z*(*t*) *−* 𝔼[*Z*(*t*)], we can assume, without loss of generality, 𝔼[*Z*(*t*)] *≡* 0 to obtain

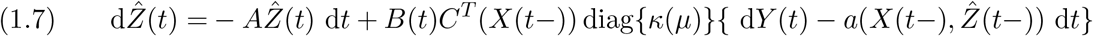

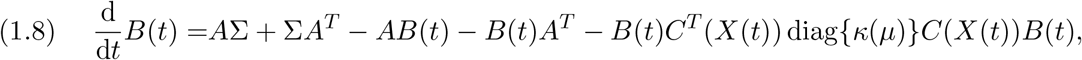

initialized in 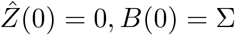, *B*(0) = ∑. We suppressed the dependency *μ* = *μ*(*X*(*t−*)) for readability. Note, that *B*(*t*) is deterministic under (C4), i.e., in the case of zeroth-order modulation.

The motivation for this Kalman-Bucy-like filter is the following, and we merely repeat here in brief the brilliant reasoning by Snyder [53, p.370-371]. Note, that it covers the case of zeroth-order modulation, as follows. The optimal linear estimator, which is to say the optimal *c*(*t*) and *h*(*t, u*), only depends on the first- and second-order information of the modulating process. Grandell [26] specified this dependence by an integral equation for *h*(*t, u*) and choosing *c*(*t*) such that the estimator is unbiased. However, the very same integral equation appears in filtering with additive noise observations instead of point process observations [36]. Hence, the optimal *h*(*t, u*) can be ”borrowed” from the Kalman-Bucy filter when assuming a mock Ornstein-Uhlenbeck process with mean 0, satisfying (C1)-(C2) and the observation model

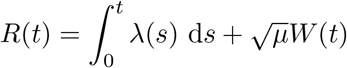

for *λ*(*t*) satisfying (C3). This is the continuous-time additive Gaussian white noise channel 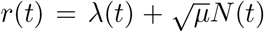 with signal-to-noise ratio *μ*^*−*1^. The Kalman-Bucy filter specifies the kernel *h*(*t, u*) in a differential form, which translates precisely to the equations Eq. (1.7)-(1.8) in the case of point process observations.

The optimal linear filter for point processes has seen some application [23, 22]. Alternatively, the Kalman filter has been used after applying a diffusion approximation [48], in the linear noise approximated case [15] or for a continuous model to begin with [17]. Consult [45] for an overview how Kalman filter and variants were applied for neural spike trains. Even without an extensive review of the literature, it seems valid to say that the alternative approach has been much more common.

We now bridge the gap from approximate state estimation to an approximate marginal description. In the case of state estimation, *Y* (*t*) is doubly stochastic with the 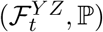 -intensity *λ*(*t*). On the contrary, we repurpose the equations for an approximate marginal description, which we regard as the core idea of this paper. Namely, we define the probability measure ℚ, such that *Y* (*t*) is self-exciting with 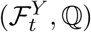-intensity

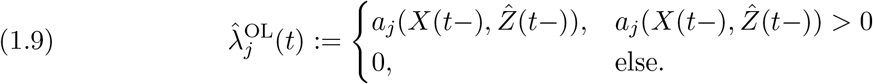

The idea is that sampling *Y* (*t*) from ℚ is simpler than from ℙ, while ℙand ℚ are, hopefully, sufficiently close. The remainder of the paper is dedicated to the question in which senseℙand ℚ are close. We provide structural results, section 2, and empirical findings, section 3. We chose here to compare two counting processes by introducing one measure for each. The reader might be more familiar with introducing a second process, denoting it by *Y* ^OL^(*t*) say, and keeping ℙ as it is. Those two perspectives are equivalent.

Before we present the results, we assume (C4) to establish a link to the (multivariate) Hawkes process with exponential kernel, introduced in the subsection 1.2. At stationarity under ℚ, Eq. (1.8) has equilibrated and the time-dependent *B*(*t*) can be replaced by the constant asymptotic value 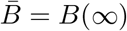. Then in integral form

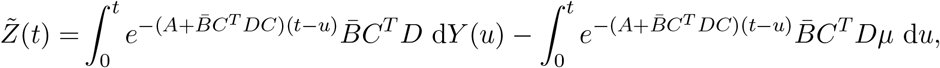

with *D* := diag^*−*1^(*μ*). In case 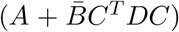 is invertible, the second term equilibrates to 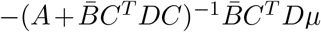. Let 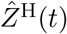 be the linear filter with both *B*(*t*) and the second term equilibrated, then 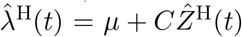 is the 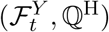 -intensity of a (multivariate) Hawkes process *Y* (*t*) with

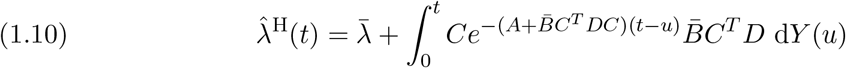

where (using the Sherman–Morrison–Woodbury formula)

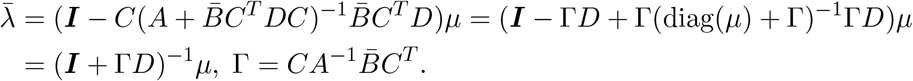

The equilibrated scenario can be made rigorous by assuming that the process is defined on the entire real axis and is at stationarity for all time *t*, provided that 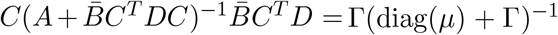 has a spectral radius strictly less than 1. In this case, the lower integral bound in Eq. (1.10) is replaced by *−∞*.

In the multivariate case, the main difference to the classical Hawkes process is the use of a matrix exponential instead of a weighted sum of one-dimensional exponential functions. The drawback of the matrix exponential is that we cannot guarantee a priori that the intensity remains positive. However, for the special case of *M* = 1 we derive a sufficient condition for a positive intensity at all times, see proposition 3.2 below. Overall, in the considered case studies, we did not encounter negative intensities. We now proceed with the time-dependent specification of *Y* (*t*) with the 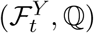 -intensity as defined in Eq. (1.7)-(1.9), which we call the *tilted* version of the Hawkes process.

## 2 Results

We have defined two different probability measures ℙ and ℚ. The measure ℙmodels the CRN in a random environment as a doubly stochastic process by means of an external process. The measure ℚprovides an approximate marginal description of the same CRN. We prove that the measure ℚ preserves the second-order properties in case the external process only modulates the zeroth-order reactions.

### 2.1 Second-order moments and spectral analysis

For the formulation of the results, we assume that

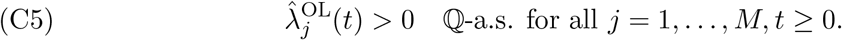

This condition implies that 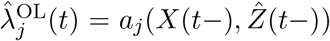 ℚ-almost surely for all *j* = 1, …, *M* and *t ≥* 0. For convenience, we assume again 𝔼[*Z*(*t*)] *≡* 0 for the external process.

#### Theorem 2.1.

*Let Y* (*t*) *be a multivariate counting process for which the* 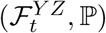 *-intensity and the external process Z*(*t*) *satisfy the standard conditions* (C1)*-*(C4), *assuming A in Eq*. (C2) *is invertible. Furthermore, let the* 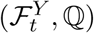 *-intensity be given by Eq*. (1.7)*-*(1.9) *and* (C4)*-*(C5). *Then for all t, s ≥* 0

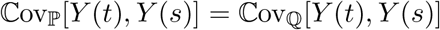

*and, with * indicating* ℙ *or* ℚ, *the expression is given by*

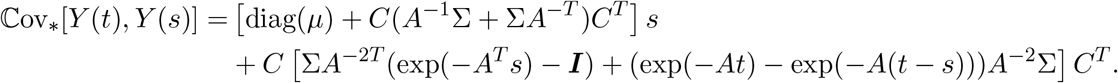

*Proof*. For the left-hand side we employ the covariance decomposition, the multivariate analogue of [11, §3.3, Eq. (3.39)], suppressing the subscript ℙ,

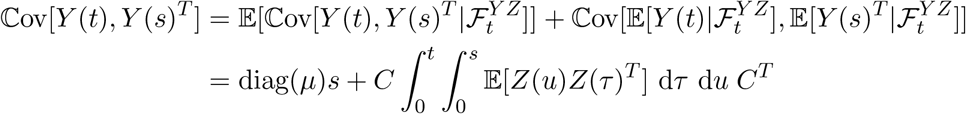

Now we split the integration domain [0, *t*] *×* [0, *s*] into three parts according to figure 1. Additionally, we change the direction of integration to the 45° diagonal (indicated by dashed lines) on which the auto/cross-covariance 𝔼[*Z*(*u*)*Z*(*τ*)^*T*^] is constant. The reason for this is the constant time lag *h* = |*τ − u*| and the assumed stationarity of the auto/cross-covariance. We get (see appendix, A)

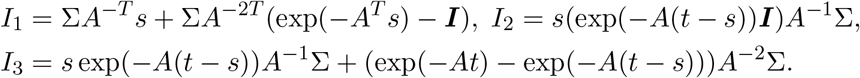

**Figure 1.**
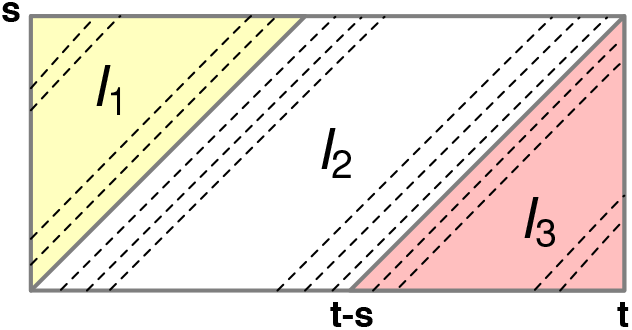
Integration domain.

In total,

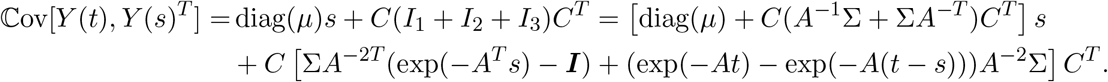

For the right-hand side we employ martingale techniques, again dropping the subscript ℚ. The martingale form of the optimal linear filter is

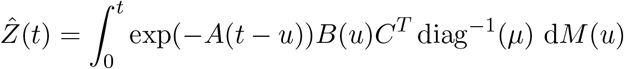

First note that the cumulative centred intensity reads

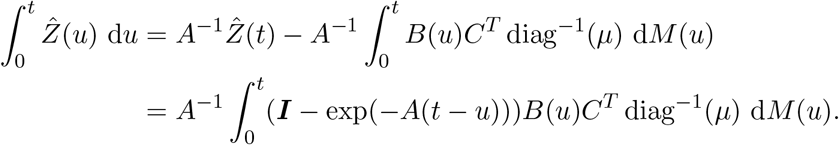

We insert this into the right-hand side as follows

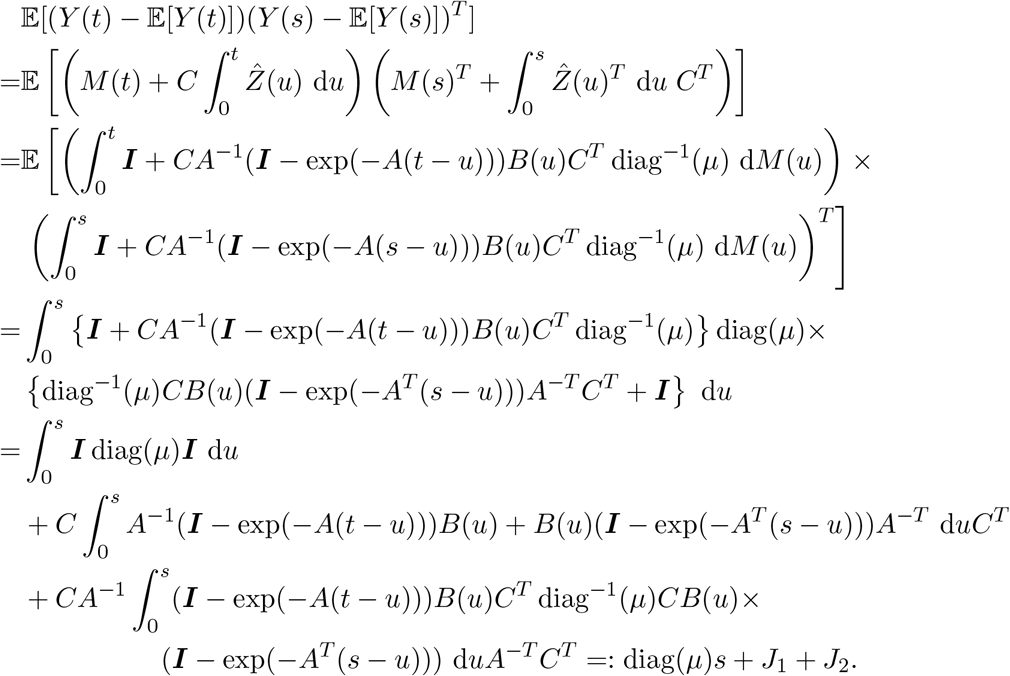

We used Eq. (1.3) in the third equality and continue

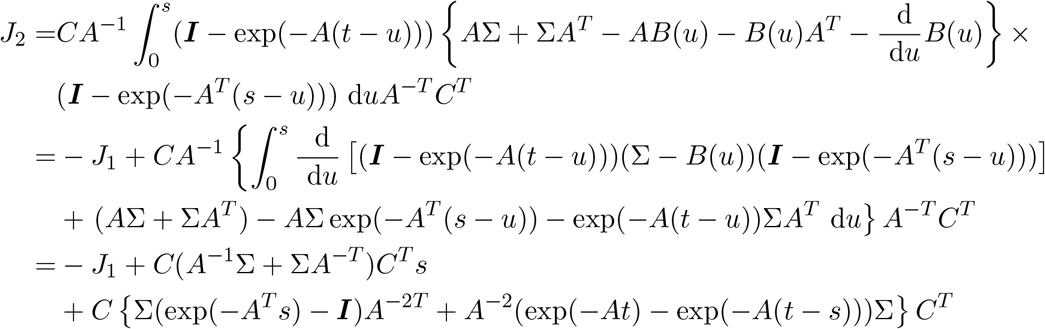

We next generalize the result to linear reaction networks in random environment, where the environment modulates the zeroth-order reactions only. To this end, we first extend the theorem 2.1 in the following lemma.

#### Lemma 2.2

*Let* [0, *t*], *u 1→ f* (*u*) *and* [0, *s*], *u 1→ g*(*u*) *be continuous and deterministic functions, then*

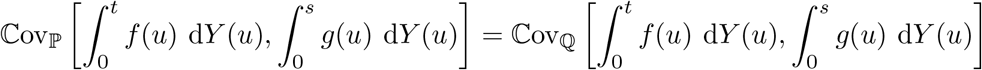

*Proof*. Define the (matrix-valued and signed) measure *C*_ℙ_ on 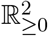 by

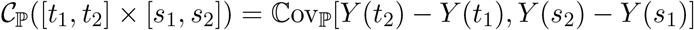

and analogously *C*_ℚ_. Then from theorem 2.1 it follows by expanding the term on the right-hand side that *C*_ℙ_ = *C*_ℚ_ on the rectangles [*t*_1_, *t*_2_] *×* [*s*_1_, *s*_2_]. Since those form a Π-system generating the Borel-sigma-algebra on 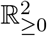, the measures *C*_ℙ_ and *C*_ℚ_ agree on 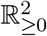. Then the claimed equality holds because the terms can be expressed as

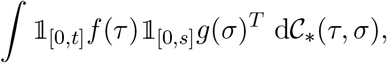

with the * indicating ℙ or ℚ respectively.

Remark: The measure *C*_*_ employed in the proof goes by the name of covariance measure in the literature, see [14, §9.5, Eq. (9.5.12)] for a principled introduction. The covariance density and complete covariance density for point processes had been introduced already at an informal basis [4] before Brémaud’s rigorous definition of point processes with stochastic intensity. The measure *C*_*\**_ has the covariance density (with respect to the Lebesgue-measure) 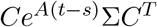, and the complete covariance density 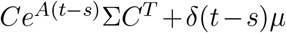. The theorem 2.1 and lemma 2.2 can thus be summarized by the observation that the doubly stochastic (multivariate) Poisson process and the (multivariate) Hawkes process have the same complete covariance density. This link was already stated in the original paper by Hawkes [30] for the univariate and the diagonalizable (orthogonal) multivariate case with exponential kernel and constant jumps at stationarity. To the best of our knowledge, the tilted version with time-dependent jumps that corresponds to the stationary doubly stochastic case, but starting at *t* = 0, is new. This offers a solution to the problem of simulating a stationary Hawkes process even when the past is not available to avoid the transient burn-in phase.

Let now *X*(*t*) = [*X*_1_(*t*), …, *X*_*n*_(*t*)]^*T*^ be the state vector of a linear CRN in a random environment, i.e., there are only first- and zeroth-order reactions. We make the assumption that the external process only modulates the zeroth-order reactions. To formalize this, we assume that the reactions R_1_, …, R_*m*_ are modulated, and the reactions R_*m*+1_, …, R_*M*_ are not modulated. Correspondingly, the stoichiometric matrix decomposes into

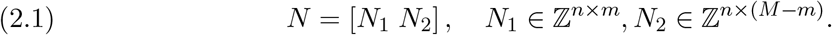

Let *Y* (*t*) ∈ ℕ^*m*^ count the occurrence of reactions R_1_, …, R_*m*_ and 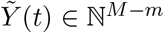 count the occurrence of reactions R_*m*+1_, …, R_*M*_. Then the state is obtained as

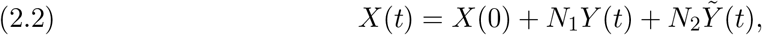

compare Eq. (1.2). We consider two probability measures ℙ and ℚfor the process *X*(*t*). Let *Z*(*t*) be an external process satisfying (C1) and (C2) with respect to ℙ. Define the filtrations

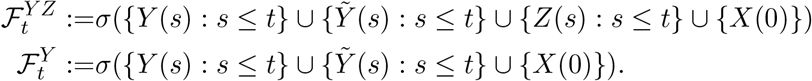

The 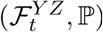 -intensity of the components *Y* (*t*) of the joint counting process [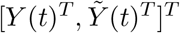 are assumed as in (C3) and (C4). The 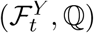 -intensity of the components *Y* (*t*) are assumed as in Eq. (1.7)-(1.9) with (C4). Furthermore, assume (C5) holds for all *j* = 1, …, *m* ℚ-almost surely. The 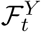 -intensity of the components 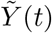 is set to be

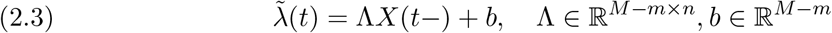

for both measures ℚ and ℙ.

#### Theorem 2.3.

*With the probability measures* ℚ *and* ℙ *as just defined and assuming X*(0) *has the same distribution for* ℚ *and* ℙ, *it holds that*

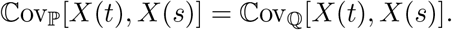

*Proof*. Without loss of generality, we assume that 𝔼[*Z*(*t*)] *≡* 0, such that 𝔼_ℙ_[d*Y* (*t*)] = 𝔼_ℚ_[d*Y* (*t*)] = *μ*. By 𝔼_*\**_, we indicate that either ℙor ℚ makes the statement correct. We decompose the differential form of Eq. (2.2) via the martingale increment 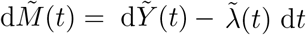

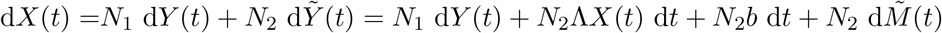

such that in integral form it holds with Γ := *N*_2_Λ

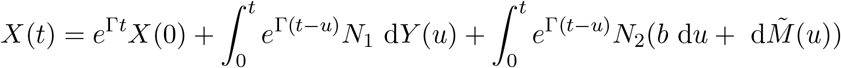

from which we conclude

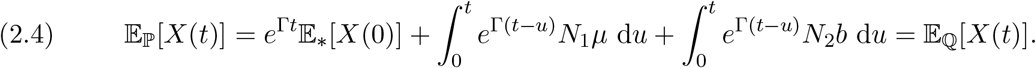

Then for the deviation of the mean Δ*X*(*t*) := *X*(*t*) *−* 𝔼_*\**_[*X*(*t*)] both with respect to ℙ and ℚ

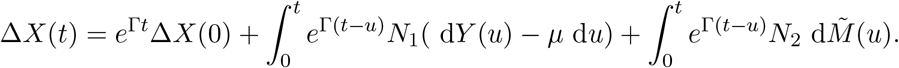

Then we obtain

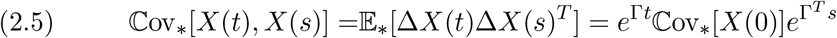

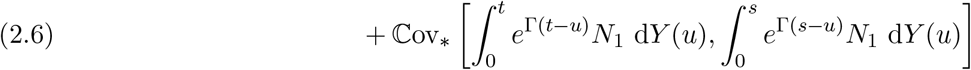

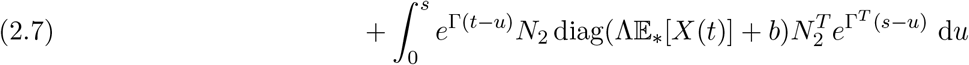

by the application of Eq. (1.6) and Eq. (1.3). All three terms agree for the measures ℙ and ℚ, the first one by assumption, the second one by the lemma 2.2 and the third one by Eq. (2.4)

We now provide the second-order result for ℙ in the frequency domain. We may drop the condition (C2) and assume a general auto/cross-spectrum *S*_*Z*_(*ω*) ∈ ℝ^*m×m*^,

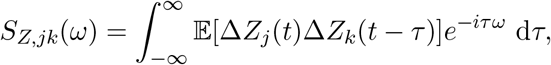

where Δ*Z*_*j*_(*t*):=*Z*_*j*_(*t*) *−* 𝔼_ℙ_[*Z*_*j*_(*t*)]. Define the probability measure 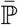, under which the intensities of *Y* (*t*) are replaced by their averages *μ*, i.e., *Y* (*t*) is a homogeneous Poisson process. Then under 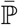, the reaction system for *X*(*t*) has propensities

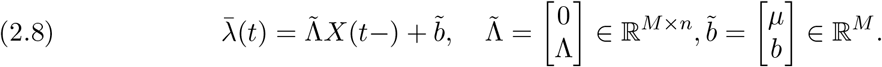

Its mean evolution equation is

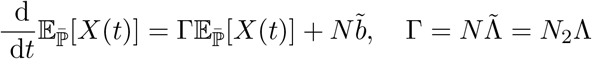

with stationary mean 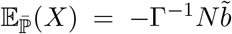. Denote by 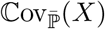 the stationary covariance matrix of the system *X*(*t*) under 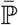. Setting up the covariance evolution equation it can be shown that 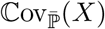satisfies

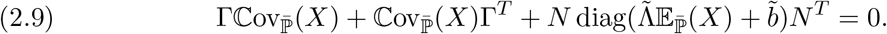

#### Proposition 2.4.

*Assuming that X*(*t*) *is the stationary process with Eq*. (2.1)*-*(2.3) *where for Y* (*t*) (C1), (C3), (C4) *hold for an external process Z*(*t*) *with auto/cross-spectrum S*_*Z*_(*ω*). *Then the auto/cross-spectrum of X*(*t*) *is*

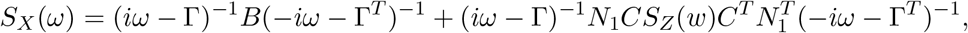

*where* 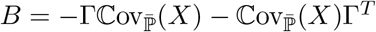.

*Proof*. We first expand the term Eq. (2.6) to the following expression by using the 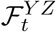 -martingale *M* (*t*)

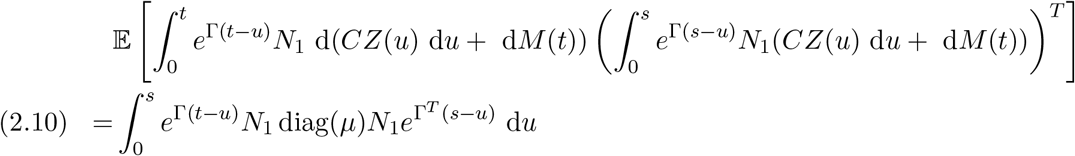

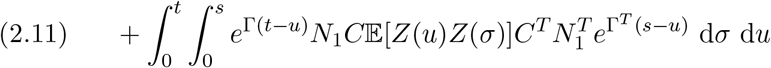

using Eq. (1.6) and Eq. (1.3). For convenience denote the term Eq. (2.11) by *L*(*t, s*) We add the terms Eq. (2.10) and Eq. (2.7) to yield

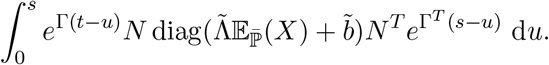

By Eq. (2.9) and using the product rule, this term evaluates to

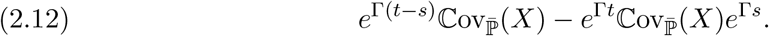

The covariance ℂov_ℙ_[*X*(*t*), *X*(*s*)] is now the sum of the terms in Eq. (2.5), Eq. (2.11) and Eq. (2.12). Letting *t* = *s* and *t → ∞*, we may decompose

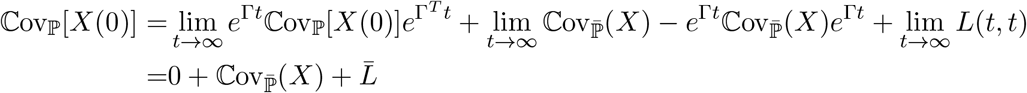

Inserting this decomposition into Eq. (2.5), we get

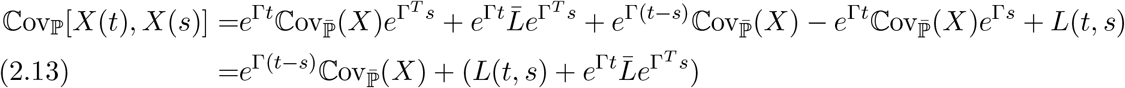

Now, in this decomposition the first term is precisely 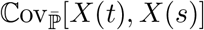. We further recognize the bracket term as the auto/cross-covariance 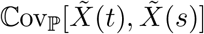of the system

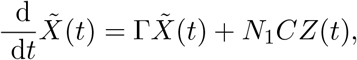

which is initialized in its stationary distribution. By the linearity of the Fourier transform, the decomposition Eq. (2.13) transfers to the spectrum as *S*_*X*_(*ω*) = *S*_1_(*ω*) + *S*_2_(*ω*). The first term yields

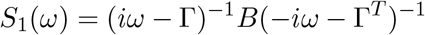

by the corresponding result on linear CRNs, see [54, Appendix 3] and [25, §4.5.6]. For the second term, we employ the transformation rule for spectra of linearly filtered stationary stochastic processes in continuous time, i.e., the continuous version of [37, §2.5 Eq. (2.94)] or the multivariate version of [38, §4.12 Eq. (4-162b)], see, e.g., [5, §7.5.1 Eq. (7.163)]. With the impulse response function 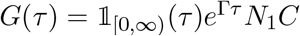 and its Fourier transform

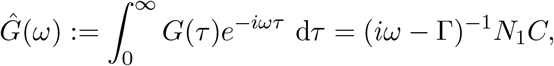

we derive the following expression

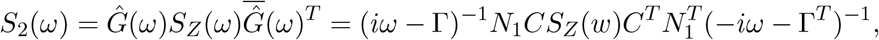

which concludes the derivation.

This proposition generalizes the corresponding result by Gupta and Khammash [28, theorem 2.1], who assigned the environment components to separate reactions and assumed them to be stochastically independent. Furthermore, the theorem generalizes the result in queuing theory, which corresponds to the birth-death with a Hawkes model for the birth events. The expressions that we provide shed light on the structure of the auto/cross-covariance, e.g., [16, Eq.(64)&(65)].

### 2.2 Characterization of the optimal linear filter

In section 1.6 we reviewed the optimal linear filter obtained from a projection method. We now present a result that characterizes it as a moment closure.

#### Theorem 2.5

*Suppose under a probability measure* ℙ, *the* 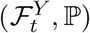*-intensity* 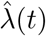*of Y* (*t*), *strictly positive in all components for all t ≥* 0, *is given by means of a predictable process V* (*t*) ∈ ℝ^*l×l*^

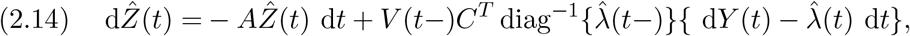

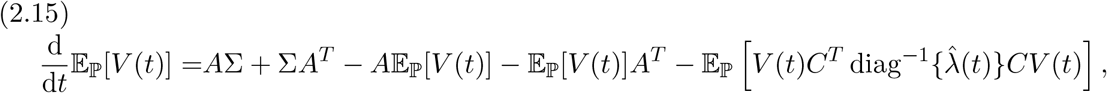

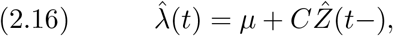

*initialized in* 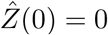, *V* (*t*) = Σ. *Then the following statements are equivalent*.

i. *The process* 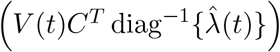 *is deterministic.*
ii. *The moment closure holds for all t≥0*

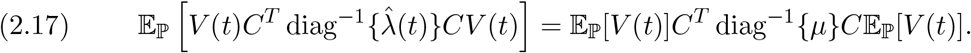

*Furthermore either one implies that* 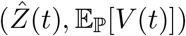 *are equal to* 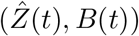*in Eq*. (1.7)*-* (1.8).

Remark: Note that Snyder’s exact filter has the form for the appropriate *F* and *G*, when using that in the evolution equation for *V*_*ik*_(*t*), the expectation of the second term is zero by Eq. (1.6). For the linear reaction networks as well as the CIR process, *F* and *G* are in the appropriate forms to match.

For the proof, we need the following generalized Cauchy-Schwarz inequality.

#### Lemma 2.6.

*Let X*_*i*_, *i* = 1, …, *m be strictly positive random variables and Y* = (*Y*_1_, …, *Y*_*m*_) *be a random vector. Then*

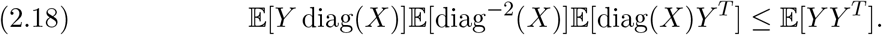

*Furthermore, equality holds if and only if for every i* = 1, …, *m there exists a deterministic scalar α*_*i*_, *such that Y*_*i*_ = *α*_*i*_*X*_*i*_.

*Proof*.

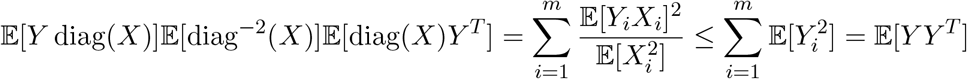

by the term-wise application of the classical Cauchy-Schwarz inequality. The additional statement holds because the sums are equal if and only if term-wise equality holds.

*Proof (theorem 2.5*). (*i*) *⇒* (*ii*). Denote 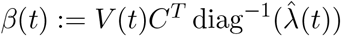. Then we compute

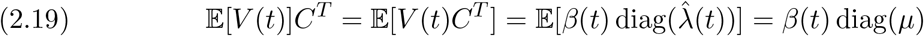

and we obtain

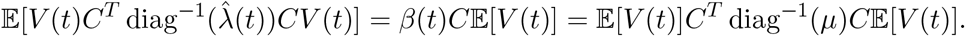

(*ii*) *⇒* (*i*). For each *k* = 1, …, *n* apply the lemma for the choice 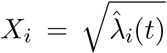 and 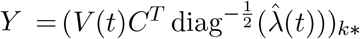 denoting the *k*-th row. Then the equality of the *k*-th diagonal entry in Eq. (2.17) reads as Eq. (2.18). Consequently, we obtain deterministic scalars *α*_*ki*_(*t*) from the lemma that satisfy

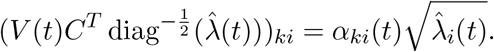

We recognize the right-hand side as the *ki*-entry of *α*(*t*) 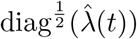. Then in matrix notation,

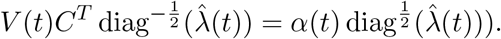

Upon multiplication with 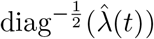 (*ii*) follows.

Let us assume that (i) and (ii) hold. The Eq. (2.19) implies *β*(*t*) = 𝔼[*V* (*t*)]*C*^*T*^ diag^*−*1^(*μ*) and together with Eq. (2.17), this yields Eq. (1.7)-(1.8) for *B*(*t*) = 𝔼[*V* (*t*)].

### 2.3 Algorithm for approximate marginal simulation

On the one hand, the joint simulation of the external process *Z*(*t*) and the reaction system *X*(*t*) provides samples from the marginal *X*(*t*). On the other hand, the 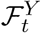-intensity offers an alternative which circumvents the co-simulation of the external process. We therefore call it the marginal simulation. Simulation with an approximate instead of Snyder’s exact filter is denoted here by approximate marginal simulation. It requires a simulation algorithm that can handle time- and history-dependent propensities. The modified next reaction algorithm by Anderson [2] proceeds by assigning each reaction counter an internal time that progresses as a unit Poisson process. Then the internal time is transformed to the global time via the inverse of the cumulative intensity. We now apply this algorithm for the simulation of a reaction system *X*(*t*) that has probability ℚ, i.e., with 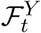intensity 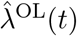 given by Eq. (1.7)-(1.9). To this end, we need to propagate the approximate state estimate 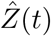and posterior covariance mock *B*(*t*) jointly between jumps via

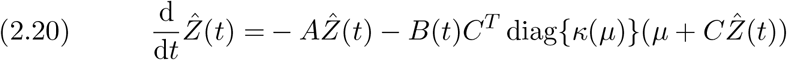

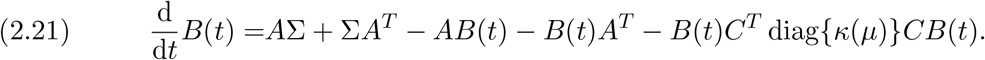

The coefficients *μ*_*j*_ and *C*_*jk*_ can be state-dependent. Additionally, each reaction R_*j*_ gets assigned another state variable Λ_*j*_(*t*), its cumulative intensity, which evolves as

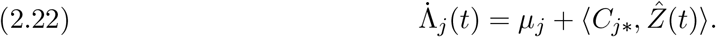

We collect them in the vector Λ(*t*) = [Λ_1_(*t*), …, Λ_*M*_ (*t*)]. For determining which reaction occurs, we need to check whether the cumulative intensity has reached the internal time difference. For this we define events, i.e., time points

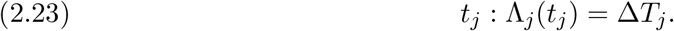

Finally, 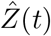 needs to be updated when the reaction R_*j*_ occurs, i.e.,

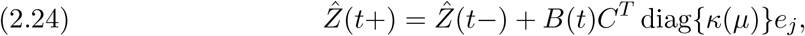

where *e*_*j*_ is the *j*-th unit vector. The complete algorithm reads as follows.

#### Algorithm 2.1 Approximate marginal simulation with the optimal linear filter

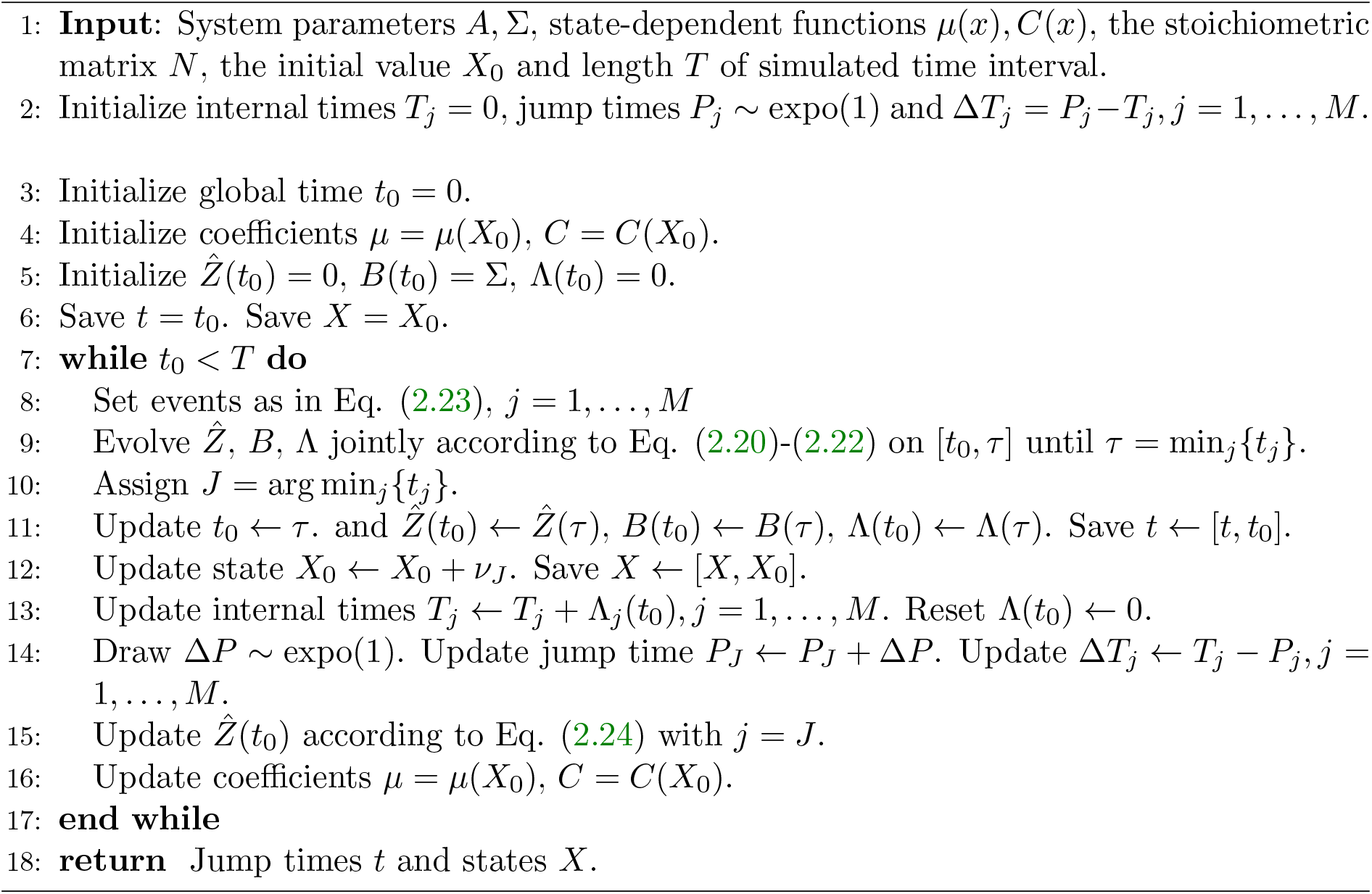

### 2.4 Independent environment components

When the environment has independent components, the line 9 of the algorithm can be simplified. Assume a diagonal structure for *A* = diag{*α*}, Σ = diag{*σ*^2^} and *C* = diag{*c*}, i.e., the environment components *Z*_1_(*t*), …, *Z*_*M*_ (*t*) are stochastically independent and *Z*_*j*_(*t*) modulates R_*j*_ via *μ*_*j*_(*X*(*t*)) + *c*_*j*_(*X*(*t*))*Z*_*j*_(*t*). In this case *B*(*t*) is also diagonal, say diag{*β*(*t*)}. We assume that *μ*_*j*_(*X*(*t*)) = 0 implies *c*_*j*_(*X*(*t*)) = 0 to avoid negative rates. We analytically solve the joint ODE system of 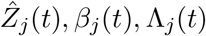 for *j* = 1, …, *M* separately. If *c*_*j*_ ≠ 0,

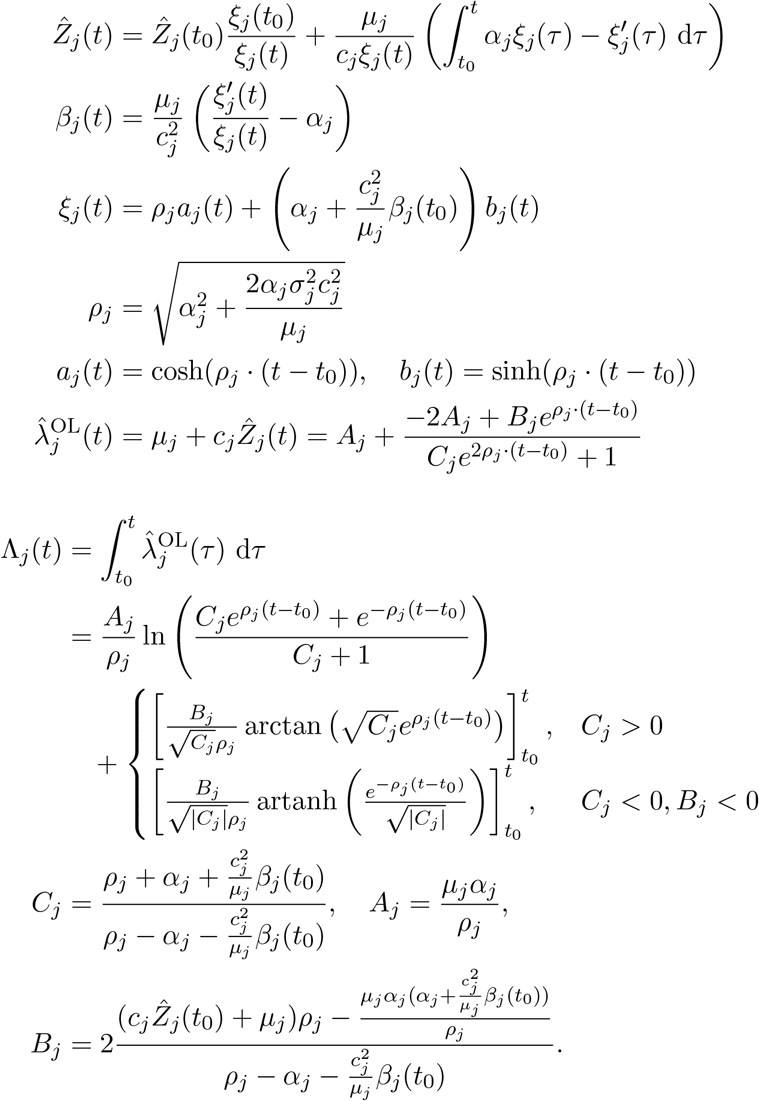

If *c*_*j*_ = 0,

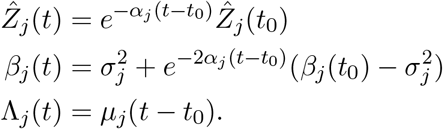

Note, that for the case *c*_*j*_ = 0 the mimics of the state estimate 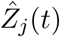and the posterior variance 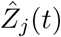 follow the prior dynamics of mean and variance. We demonstrate the use of this filter in the second case study when we investigate a two-stage gene expression in uncorrelated random environment.

## 3 Case Studies

We now illustrate the strengths and limitations of the equilibrated optimal linear (Hawkes) model, Eq. (1.10), and the tilted Hawkes model, Eq. (1.7)-(1.9), in three case studies. Whether ℚ approximates ℙ well guides our experiments.

### 3.1 Promoter-mediated transcription

The transcription of mRNA is a process that is shaped by the cellular context. The number of polymerases, fluctuating transcription factor concentrations, noncognate binding and chromatin remodelling are some of the factors that determine the rate of transcription. Often these influences are abstracted into a low number of discrete promoter states that determine the transcription rate. The effect of switching between an active and an inactive promoter state is often equated with transcriptional bursting. Suppose there are *N* transcription sites with the same promoter structure. At each site, the promoter can be in one of *k* states. The switching between the states occurs according to a Markov jump process with transition rates *c*_*ij*_ for the *i*-th to the *j*-th state. When being in the *i*-th state, the transcription rate at which mRNA are produced is *r*_*i*_. We assume the transcription sites have the same stochastic properties, i.e., the same *c*_*ij*_ and *r*_*i*_ and are stochastically independent. Assume further that mRNA decays with a rate *γ*. Then the total number of mRNA *X*(*t*) from all transcription site is a birth-death process whose birth rate is modulated by a conversion process *Z*(*t*) = [*Z*_1_(*t*), …, *Z*_*k*_(*t*)] where *Z*_*i*_(*t*) counts the number of transcription sites that are in state *i* at time *t*. The conditions (C3) and (C4) are satisfied. We assume (C1) to hold. Moreover, the conversion process is a closed system with only monomolecular (conversion) reactions, in particular it is linear and hence (C2) is satisfied. Its stationary distribution is the multinomial distribution with parameters *p*_1_, …, *p*_*k*_ and *N*. The system parameters are derived to be

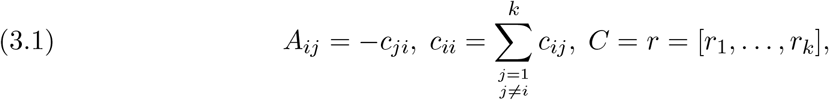

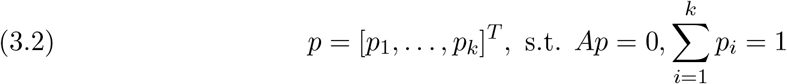

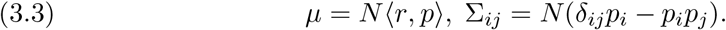

The multinomial stationary distribution implied the form of the covariance matrix, Eq. (3.3). According to Jahnke and Huisinga [32] *p* is uniquely determined by the condition in Eq. (3.2) if *A* is irreducible.

We consider the equilibrated optimal linear (Hawkes) approximation of this system, i.e., 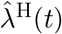as in Eq. (1.10). For this system, we provide the equilibrium value of the ODE for 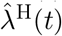between jumps, from which we compute 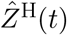 that is shown to be positive. For the derivations, we require *C* ∈ ℝ^1*×l*^and *A* to be invertible. For the conversion process, this is not the case, since **1***A* = 0, but by truncation to *l* = *k −* 1 we can enforce it, see the appendix B.1.

The equilibrium 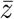 of the linear dynamics

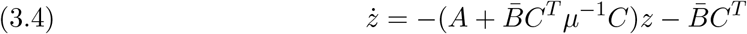

between jumps is evaluated by the Sherman-Morrison formula as

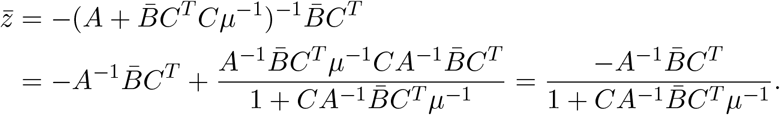

We assume that 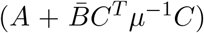 has only eigenvalues with positive real part. Then the equilibrium is stable and the equilibrium value 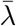 of 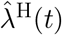between jumps is

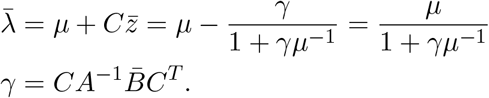

Furthermore, jumps of 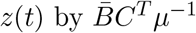 increase 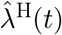by

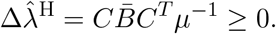

#### Proposition 3.1

*The term* 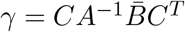 *is the positive root of*

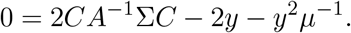

*Proof*. First, we note that 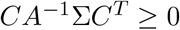, since

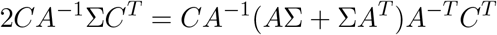

and by condition (C2) *A*Σ + Σ*A*^*T*^ is positive semidefinite. Since *B*(0) = Σ, we have

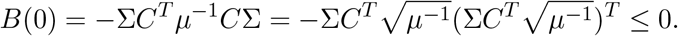

Then by [10, Corollary 3] *B*(*t*) is monotonically non-increasing. By the property of complete observability, *B*(*t*) converges monotonically to the equilibrium 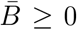. Define *x*(*t*) := *CA*^*−*1^*B*(*t*)*A*^*−T*^ *C*^*T*^. Then by the monotonicity of *B*(*t*), it holds that 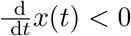. With minor algebraic manipulation, we express

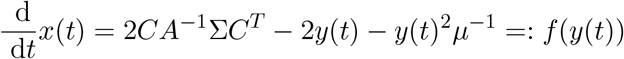

in terms of *y*(*t*) = *CA*^*−*1^*B*(*t*)*C*^*T*^. Then necessarily, at equilibrium of *x*(*t*) and *y*(*t*), it holds 0 = 2*CA*^*−*1^σ*C*^*T*^ *−* 2*γ − γ*^2^*μ*^*−*1^. The positive and the negative solution *γ*_1_ and *γ*_2_ split the real line into three intervals (*−∞, γ*_2_), (*γ*_2_, *γ*_1_), (*γ*_1_, *∞*) on which *f* (*y*) is negative, positive, negative. The initial value *y*(0) = *CA*^*−*1^Σ*C*^*T*^ is positive and *f* (*y*(0)) *<* 0, hence *y*(0) ∈ (*γ*_1_, *∞*). But since 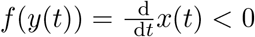for all *t ≥* 0, the solution *y*(*t*) ∈ (*γ*_1_, *∞*) for all *t ≥* 0. Hence, *γ* is the positive root *γ*_1_.

Then immediately we obtain

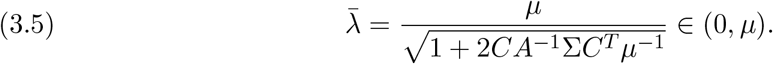

For the condition (C5) to hold we provide the following sufficient criterion which guarantees that 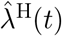stays positive at all times.

#### Proposition 3.2.

*Suppose that z*_1_, …, *z*_*l*_ *form a basis of real eigenvectors of* 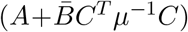*for the positive eigenvalues ν*_1_, …, *ν*_*l*_, *such that Cz*_*i*_ *>* 0 *for all i* = 1, …, *l. Let L be the matrix whose columns are the eigenvectors z*_1_, …, *z*_*l*_ *and assume that*

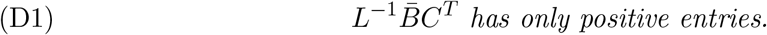

*Then* 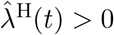*for all t ≥* 0.

*Proof*. We consider the half-space Λ_+_ := {*z*; *Cz* + *μ >* 0} with boundary {*z*; *Cz* + *μ* = 0} as well as the inward normal vector *C*, and the cone 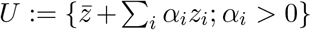. By Eq. (3.5), 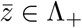 and hence *Cz*_*i*_ *>* 0 implies *U* ⊆ Λ_+_. By assumption, there exist positive *β*_*i*_, *i* = 1, …, *l* such that 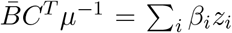 Consequently, *f* (*U*) ⊆ *U* for the map 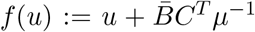, which means that jumps of the trajectory 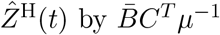remain in *U*. The initial value 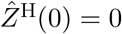is in *U* because

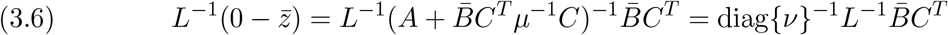

has only positive entries. Finally, we verify that the dynamics in Eq. (3.4) leaves *U* invariant. Let *z* be any boundary point of *U*, i.e. 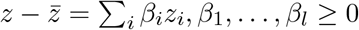 and there exists *i* such that *β*_*i*_ = 0. Let *η* be any corresponding inward normal vector, in particular, ⟨*z*_*i*_, *η*⟩ = 0 for all *i* = 1, …, *l*. Then

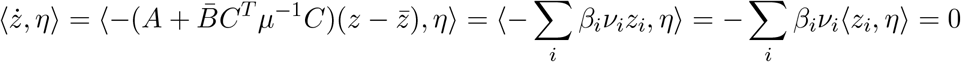

As a remark, we note that the condition D1 is equivalent to

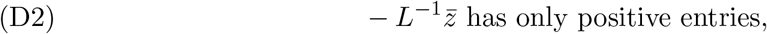

by Eq. (3.6). Geometrically, the condition corresponds to requiring that the ray starting from 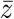 and going through 0 lies within the cone *U*, defined in the proof.

Returning to the case study of promoter-mediated transcription, we first looked at a model which unidirectionally transitions through an active, an inactive and a refractory state, see figure 2A. We chose equal transition rates, hence the promoter uniformly spends time in either state. Assuming leaky transcription, we set the transcription rate for the inactive state to a small leakage value, the rate for the active state to the highest and for the refractory state to an intermediate value. Furthermore, we considered four independent copy of the promoter. We numerically verified the condition (D1) for the considered choice of parameters. We looked at the stationary distribution of the mRNA, figure 2B to compare the exact system with the equilibrated optimal linear (Hawkes) model, Eq. (1.10). From theorem 2.3 we know that the variances coincide. However, also the shape is essentially captured. When looking at the true and approximated state estimate (fig 2C) for a mRNA trajectory (fig 2D), both also agree to a large extent. Even though the equilibrated optimal linear (Hawkes) model does not capture the base level, the state estimate seems to allocate the regime where the linearization fits well for a large amount of time.

**Figure 2.**
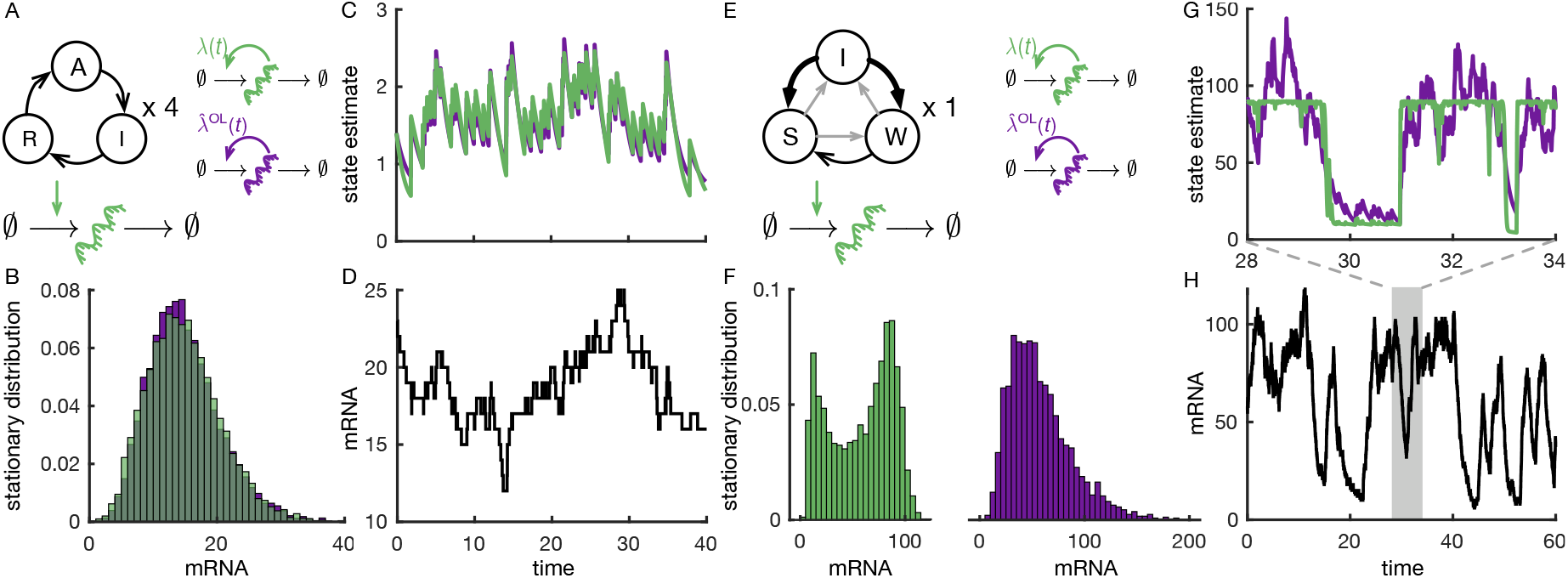
Promoter-mediated transcription. A. Transcription model with promoter states modelled by a discrete Markov process. The promoter model transitions through an active, an inactive and a refractory state with all three transition rates being equal to 0.2. Each state summarizes at what rate transcription is initiated. Assuming a leakage of 0.01, we chose exemplarily for the transcription rates 1.01 (A), 0.01 (I), 0.06 (R). Furthermore, we considered four independent copies corresponding to different transcription sites in the cell. We compared the true system (green) and the equilibrated optimal linear (Hawkes) approximation (purple). B. Histogram of the stationary distribution for the exact simulations, using Gillespie (green) and the approximate marginal simulations, using Anderson’s modified next reaction method (purple). C. State estimate of the effective (posterior mean) transcription rate obtained as an average over the posterior probabilities of being in either promoter state. Green uses the true posterior probabilities, purple the equilibrated optimal linear (Hawkes) model, Eq. (1.10). D. The mRNA trajectory for which the state estimate was computed. It was simulated, using the Gillespie algorithm. E. A different promoter model is used compared to A with an inactive, a weak and a strong state. The arrow thickness indicates the transition strength. Transition and transcription rates were c_WI_ = c_SW_ = c_SI_ = 0.2, c_WS_ = 0.4, c_IS_ = c_IW_ = 2, λ_S_ = 90, λ_W_ = 10, λ_I_ = 0 (borrowed from [59]). We considered only a single transcription site. F. Histogram as in B. The exact bimodal distribution is not captured by the equilibrated optimal linear (Hawkes) model, even though the variances agree. G. State estimates as in C for the time interval [28, 34]. H. Sample trajectory corresponding to G, simulated using Gillespie. The degradation rate of mRNA was chosen 0.1 in model A and E.

We now turn towards a promoter model, which has more extreme properties than the previous one. The new model allows transitions in all directions, where two states, a strong and a weak one, are dominantly allocated, see figure 2E. The major changes are: (i) an increase of the transcription rates compared to the transition rates, (ii) only one transcription site as opposed to four copies. Again, we numerically verified the condition (D1). Then we see that the true bimodal distribution is not captured by the equilibrated optimal linear (Hawkes) model, even though the variance is captured correctly (fig 2F). Looking at the exact and the approximate state estimates (fig 2G and H) we see that the very sharp transitions are not well captured by the equilibrated optimal linear (Hawkes) model. These non-linearities are out of reach for the linearization. The sample trajectory (fig 2H) was simulated with the exact system (Doob-Gillespie). An mRNA trajectory simulated by the approximate marginal simulation is shown in the appendix, figure 7. It differs from the exact system in underestimating the time spent in the regime of 80 to 100 mRNA copies, compare also the histograms in figure 2F.

We hypothesized that both a relaxation of (i) and (ii) could lead to a better agreement with the linearization. As a necessary condition, the bimodal distribution must transform to a unimodal one. To relax (i) we multiplied the transition rates by a factor *a >* 1, and to relax (ii) we varied *N*. In the appendix, figure 8 we depict the corresponding histograms for *a* = 1, …, 5 and *N* = 1, …, 5, which provides evidence that the necessary condition is met.

If in the case of relaxing (i) the timescale of the promoter approaches the timescale of the transcription then the knowledge about the state is more vague, hence the domain of state estimate values that are allocated narrows. On this domain, the linearization can have a good accuracy for a larger proportion. The extreme case would be a much faster promoter timescale than the transcription timescale, for which the state estimate distribution would narrow to a peak at the mean value. If in the case of relaxing (ii) we increase the number of copies, the prior distribution of intensity values becomes unimodal, in the sense that the distribution of 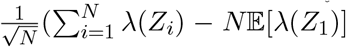 approaches a Gaussian for the i.i.d. promoter copies *Z*_*i*_ distributed as the stationary prior distribution of one promoter. Let us assume the extreme case of perfect state estimation, with the distribution of state estimate values equal to the prior distribution of intensity values. This distribution would resemble more the distribution of intensity value a Hawkes model can achieve, see [50]. With imperfect state estimation, the linearization is even improved by the above argument. In summary, we conjecture that both with a lower frequency of point observations and with a more Gaussian shape of the prior distribution of intensity values, the accuracy of the equilibrated optimal linear (Hawkes) model can increase.

### 3.2 Gene expression in a random environment

In the previous case study we considered a modulated zeroth-order reaction by a correlated environment. Now we illustrate the simulation algorithm for an environment with independent components, section 2.4, that corresponds to the tilted Hawkes model. Each component modulates one of the four reactions of the following gene expression model, see figure 3A.

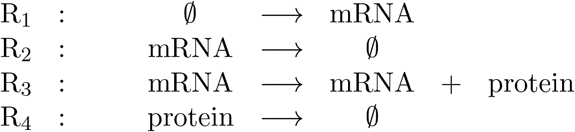

**Figure 3.**
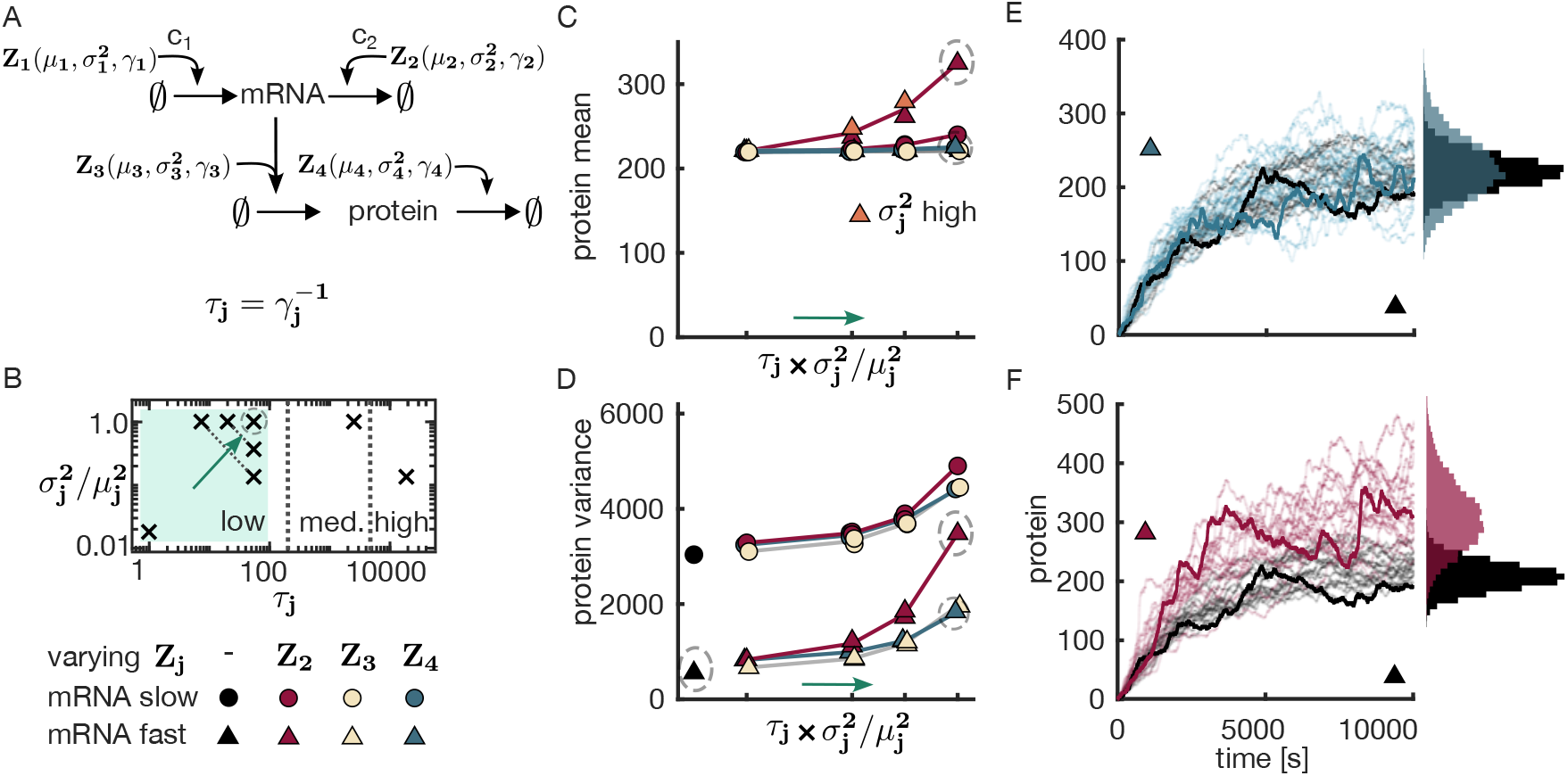
Gene expression model in a fast random environment. A. Four independent environment components modulate the gene expression model. Each component Z_j_ is characterized by its mean μ_j_, its variance 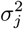 and its autocorrelation decay γ_j_. The inverse of γ_j_ is the autocorrelation time τ_j_. The birth and death reactions of the mRNA can be multiplied by factors c_1_ and c_2_. B. The setup of the simulation study -We simulated trajectories of the model in A with the approximate marginal simulation algorithm for the tilted Hawkes approximation. For each component Z_2_, Z_3_ and Z_4_, we varied 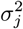and τ_j_. The parameter pairs 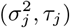 for which we simulated trajectories with are marked by crosses in the logarithmic 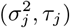-plane. For comparison between the three components, we varied 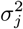relative to 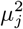. Along the decreasing diagonals, the product 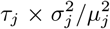 is constant. We distinguished the low autocorrelation regime from the medium and high regime. A correlation time of 50 − 500s (upper end of low regime to lower end of the medium regime) was the correlation time of the mRNA. We conducted simulations for a slow (c_1_ = c_2_ = 0.1) and fast (c_1_ = c_2_ = 1) mRNA timescale. The black circle and triangle symbolizes the reference model, in which all environment components are replaced by their constant mean values. C and D. The mean and the variance of the protein at time point t = 10000, estimated from 10000 sampled trajectories, is shown for the different conditions. The ticks of the x-axis correspond to the four diagonals of the low regime in B. Accordingly, the second and third tick combine two values which overlap, sometimes to an extent that only one is visible. For Z_2_ we indicated the pair with the larger 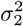 by orange (C). Lines were added and different colours for the same tick were shifted horizontally to improve readability. The stationary variance of the homogeneous model was added for reference in black. E and F. Sample protein trajectories of the reference model were plotted in black to contrast the embedded model with the largest variance and correlation for fast mRNA timescale and Z_4_ (E) and Z_2_ (F). Histograms of time t = 10000 were added. Circled marks in B, C and D indicate that for these parameter choice trajectories are shown in E and F. The components that were not varied were fixed at 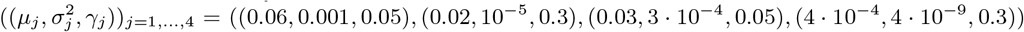. In the shaded low regime of B parameters were 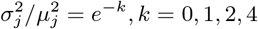and 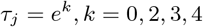.

In particular, compared to the previous case study, we also incorporate modulation of the first-order reactions in this example. We then asked the question of how the characteristics of the environment components impact the statistical properties of the protein trajectories. When all environment components are set to their constant mean level, we recover the unembedded gene expression model, which we call homogeneous or reference model. As relevant characteristics of the environment, we considered the variance and the autocorrelation time of the individual environment components. We varied these characteristics for the first-order modulators *Z*_2_, *Z*_3_ and *Z*_4_ one at a time, while leaving the remaining ones fixed. Figure 3B depicts the setup of the simulation study. For the variance 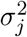, we chose 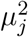 as an upper bound for the considered domain. This bound is motivated by the CIR process, which guarantees the strict positivity of the trajectory only for 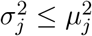. For the autocorrelation time 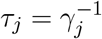, we distinguished three regions. The region for which the timescale of changes in the environment is of the order of the mRNA timescale or faster (low autocorrelation), the region where the environment timescale is shared with the protein timescale (medium autocorrelation) and the slower-than-protein timescale (high autocorrelation). Additionally, we wanted to investigate how the timescale differences between the mRNA level and the protein level in the reference model affects the results. For this purpose, we separately investigated a fast mRNA timescale (*Z*_1_ and *Z*_2_ multiplied by *c*_1_ = *c*_2_ = 1) and a slow mRNA timescale (*Z*_1_ and *Z*_2_ multiplied by *c*_1_ = *c*_2_ = 0.1). This choice keeps the mean of the protein level the same for the homogeneous case, but brings the mRNA and protein timescales closer to each other in the latter case. The gene expression model is a cascade of two birth-death processes. We thus were interested to understand what was different between changing the modulation at the mRNA level (R_2_) compared to the protein level (R_3_ and R_4_). Also, we expected to see differences between the modulation of the birth reaction (R_3_) and the death reactions (R_2_ and R_4_).

We first investigated the regime of low environment correlation time, at most of the order of the mRNA level. We saw no effect in the stationary mean of the protein (figure 3C) for varying the modulation at the protein level (R_3_ and R_4_). For a variation at the mRNA level (R_2_), the stationary mean increased when we increased 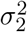or *τ*_2_. This effect was stronger for the fast mRNA timescale and for larger 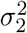 when the product 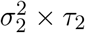was fixed. The autocorrelation showed no substantial change in either case, see appendix, figure 9. In contrast, we saw a noticeable effect on the stationary variance of the protein in all conditions, see figure 3D. The variance of the reference system was calculated as [54]

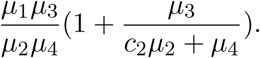

It is composed of a Poissonian and an additional term. The Poissonian term accounts for the intrinsic variance at the protein level, while the additional term captures the impact of the mRNA level. The latter is dominated by the mRNA timescale *c*_2_*μ*_2_, which is larger than *μ*_4_. As a consequence, we get a significantly higher variance level for the slow mRNA timescale, which is attributed to the effect of the mRNA level on the protein noise. On top of these two reference values of the variance, the modulating environment further increases the variance. We found that the effect was the strongest for a modulation at the mRNA level by R_2_, while R_3_ and R_4_ showed a similar effect. Figure 3E illustrates the broader distribution compared to the reference model for the modulation by *Z*_4_ with maximal 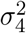 and *τ*_4_. Can the increase be attributing to either an increased environment autocorrelation time *τ*_*j*_ or environment variance 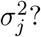 We obtained Monte Carlo estimates of the variance at time *t* = 10000 for different pairs of autocorrelation time and variance, see figure 3B. (maybe shift next sentence to figure caption.) Their product is constant along descending diagonals in the logarithmic plane of *τ*_*j*_ and 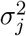. Interestingly, the simulation studies seem to indicate that their product majorly determines the variance level. This observation is in line with the increase of the variance for the zero-order modulation.

An exception to this observation is the condition with a fast mRNA timescale and varying R_2_ modulation, which shows a steeper increase in the variance compared to the other conditions (fig 3D). Also for the slow mRNA timescale the R_2_ modulation shows a deviation compared to R_3_ and R_4_ in the condition with the maximal 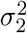and the highest *τ*_2_. Note the parallel to the protein mean in figure 3C. Sample trajectories for the modulation of R_2_ with the largest values of *τ*_2_ and 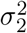 and for the fast mRNA timescale are depicted in 3F. They are contrasted with trajectories from the reference system. We observe the increase in both the variance and the mean. In summary, we found that in the regime of low environment autocorrelation (the order of the mRNA at most), there is an effect on the variance for all conditions, which becomes significant only when *τ*_*j*_ approaches the autocorrelation time of the reference mRNA level and 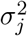approaches the maximal value. For the modulation of the mRNA level, R_2_ there was a higher increase compared to R_3_ and R_4_, whose increase majorly depended on the product of *τ*_*j*_ and 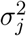. This additional contribution is likely associated with the increased protein mean for this condition. The trajectories look qualitatively the same as for the reference model.

We secondly investigated the regimes of medium and high environment autocorrelation. We were interested whether our finding on the variance for the low regime still holds, or whether factors other than the product of *τ*_*j*_ and 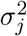 determine the variance. The exception of R_2_ modulation hints at other contributing factors. For this purpose, we chose two pairs of autocorrelation time and variance, which have the same product, figure 4A. One pair utilizes the maximal variance combined with a medium environment correlation time which is of the order of the reference protein timescale. The other pair utilizes an autocorrelation larger than the autocorrelation time of the protein, combined with a medium value for the variance. In the protein mean, we saw a different behaviour for the modulation of death reactions compared to the birth reaction, figure 4B. For R_2_, we observed largely increased values in the means for the first (correlation time, variance)-pair compared to the second. For R_4_, we observed an increase comparable with the increase for R_2_ in the low regime under fast mRNA timescale. In the variance, we noticed different values of both pairs for all reactions, figure 4C and also the autocorrelation exhibits a noticeable difference 4D. Before we investigate these differences further, we first note, that for the mean, it was to be expected that there is no effect when the birth reaction R_3_ is modulated. The mean equations read

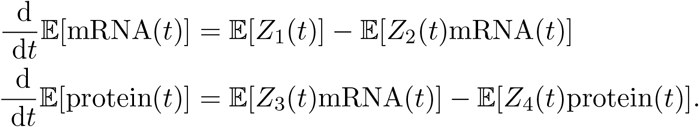

**Figure 4.**
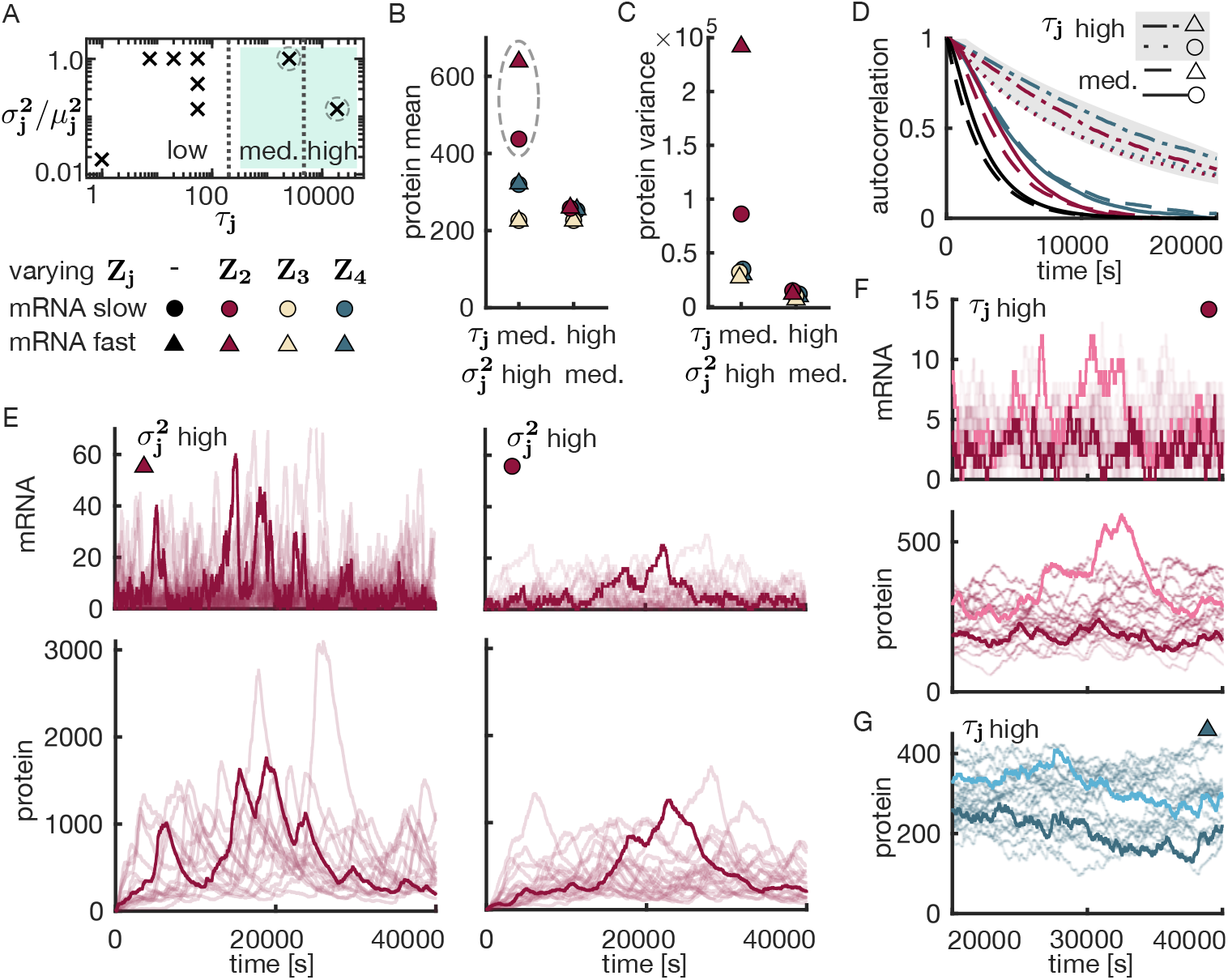
Gene expression model in a slow random environment. A. As fig 3B, but with focus on the medium and high correlation time. The mark in the medium regime is the timescale τ_j_ = μ_4_ of the reference model. We simulated trajectories of the model in fig 3A with the approximate marginal simulation algorithm (tilted Hawkes approximation) for the two marked parameter pairs. Circled marks indicate that E depicts corresponding trajectories. B and C. The protein mean and variance at time t = 40000 estimated from 10000 sampled trajectories for the different conditions. Different colours for the same tick were shifted horizontally to improve readability. D. The protein autocorrelation as a function of the lag. Black trajectories indicate the analytically computed autocorrelation of the reference model. Coloured curves indicate the correlation coefficient of protein trajectories for the embedded model at different conditions of varying Z_2_ and Z_4_, estimated from 10000 sampled trajectories. For reference model and medium τ_j_, dashed = fast mRNA timescale, solid = slow mRNA timescale. The high correlation regime is shaded by grey (dashed-dotted = fast mRNA timescale, dotted = slow mRNA timescale). E. Sample trajectories of the mRNA and the protein for the maximal 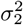 value and medium (protein-like) correlation time τ_2_. The left mRNA-protein pair depicts the fast mRNA timescale and the right one depicts the slow mRNA timescale. F. Sample trajectories of the mRNA and the protein for the pair of a high (more-than-protein) correlation time τ_2_ and a medium 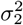 value with slow mRNA timescale. Trajectories are depicted from t = 20000 after stationarity has been reached. G. Sample trajectories of the protein for the pair of a high (more-than-protein) correlation time τ_4_ and a medium 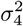value with a fast mRNA timescale, at stationarity as in F. In the shaded medium and high regime of A parameters were 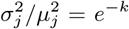and τ_j_ = 2500 · e^k^, k = 0, 2.

**Figure 5.**
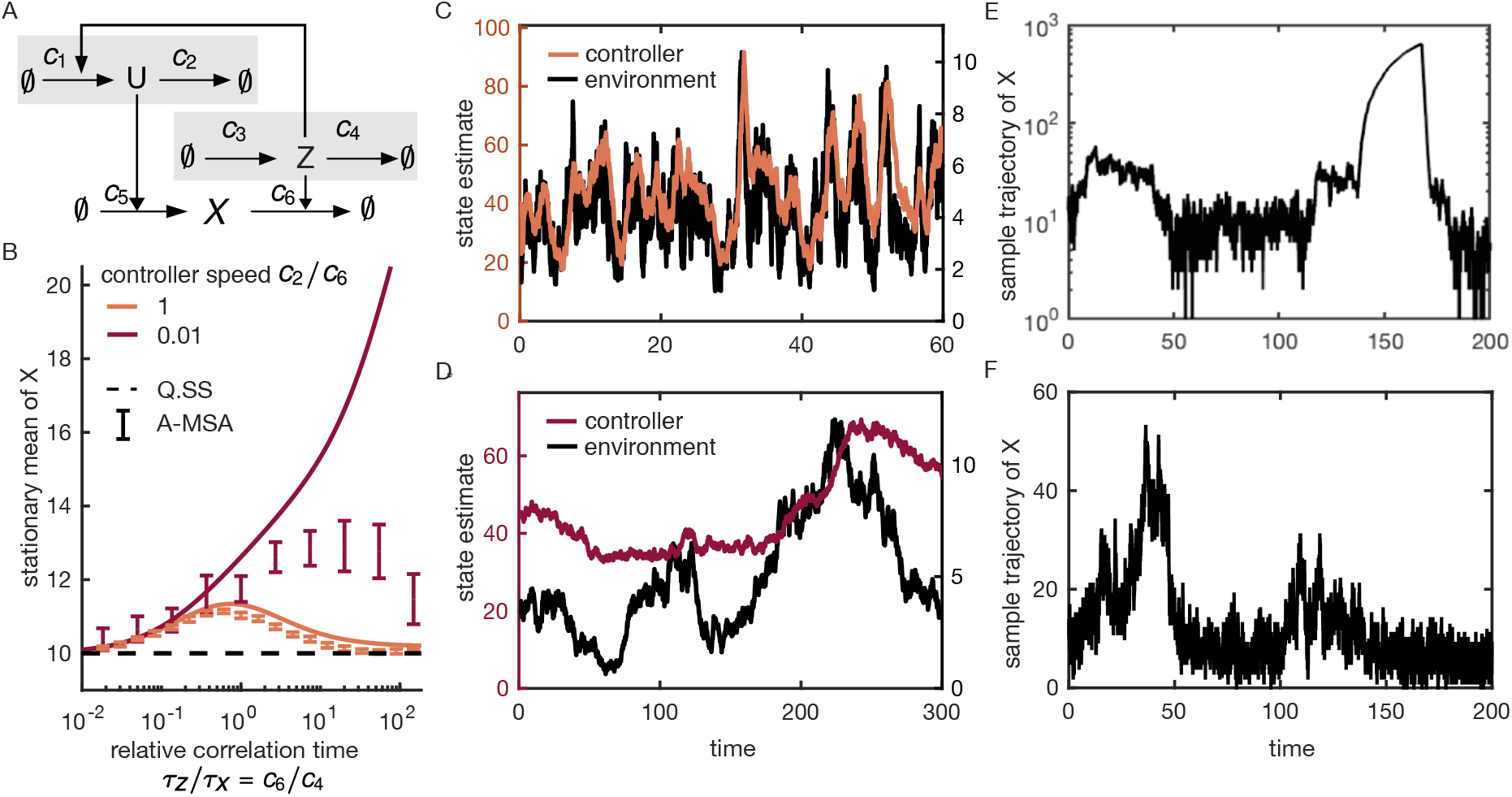
Modulation of first-order reaction. A. A birth-death process X is embedded into a correlated environment (U, Z). The controller U senses the environment component Z which modulates the death rate of X, and it regulates the birth rate of X accordingly to achieve the set point goal of keeping X near 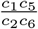. B. The stationary mean is plotted for two controller speeds as a function of the environment speed, dictated by the relative correlation time c_6_/c_4_. The environment becomes slower from left to right. As a reference, the set point (quasi-steady state) is added. The stationary mean serves as a proxy to assess whether the control goal is achieved. The error bars indicate the stationary mean of the tilted Hawkes model (approximate marginal simulation algorithm). For each of the 20 replicates it was computed as the temporal average of a trajectory of 55, 000 (100, 000) transitions, truncated by a burn in period of the first 5000 (50, 000) transitions for the fast (slow) controller. The 95% bootstrap confidence interval for the mean, using 1000 bootstrap samples of the 20 replicates, is shown. C. Sample trajectory for the approximate state estimates of the controller U and the environment component Z in the system with a slow controller. D. As in C for a slow controller. E. Sample trajectory of X for the slow controller (Doob-Gillespie). F. Sample trajectory of X for the slow controller (approximate marginal simulation). Parameters were c_1_/c_2_ = 10, c_3_/c_4_ = 4, c_5_ = c_6_ = 1 and c_2_ = 1, c_6_/c_4_ = 0.6 (C), c_2_ = 0.01, c_6_/c_4_ = 54.6 (D-F).

**Figure 6.**
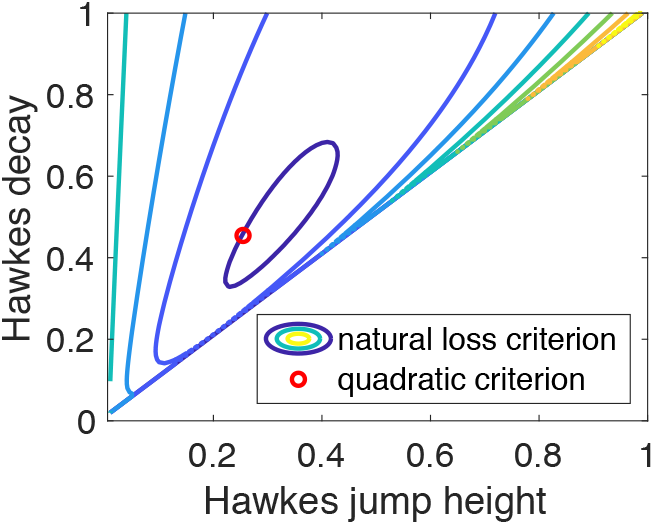
Optimal Hawkes parameters. Exemplarily, we chose a random telegraph modulated counting process model. Consider the task of finding the optimal one-dimensional Hawkes model 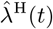 with parameters as in Eq. (1.4). Level sets indicate the natural loss criterion 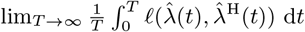 in the (β, α)-plane. The red circle locates the optimum with respect to the quadratic criterion. The stationary mean μα/β of the Hawkes intensity is equal for both criteria.

Since the environment component *Z*_3_(*t*) is independent of mRNA(*t*), the first term in the second line equals *μ*_3_𝔼[mRNA(*t*)]. Hence, the mean equations do not depend on 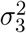and *γ*_3_. Consequently, the mean protein level is expected to be the same as in the low regime. Now, we turn to the more interesting modulation of the death reactions. For the modulation of R_4_, we notice a difference in the protein means. From looking at sample trajectories, see appendix, figure 10A, we conclude that periods of low environment value *Z*_4_ can cause excursions, meaning periods of strong deviations from the mean, in the protein. We further observe that the mean does not depend on the relative timescales of the mRNA and protein. A way to explain this is by arguing that the protein mean depends linearly on the average protein birth rate. The protein birth on average is majorly affected by the mRNA mean. The mRNA however is the same for both the fast and the slow mRNA timescale. Conversely, this means that when we see a difference in the protein mean for the two timescales of the mRNA, this might be attributed to different mean mRNA levels. And indeed, for the modulation of R_2_, we see a significant difference between both timescales (figure 4B). We confirmed that this difference can be attributed to a difference in the mean levels of the mRNA, see appendix, figure 10B. By looking at sample trajectories in figure 4E, we see excursions at the mRNA level. By the same argument as for the R_4_ modulation at the protein level, these can be attributed to phases of low *Z*_2_. To explain the difference between the two mRNA timescales, we see that the excursions are heavier for the fast mRNA timescale (left-hand side of 4E) than for the slow one (right-hand side of 4E). This can be explained by a higher average mRNA birth rate *c*_1_*μ*_1_ for *c*_1_ = 1 compared to *c*_1_ = 0.1, while the length of the excursions depends mostly on *τ*_2_, which is the same for both cases. Looking for other quantities that are equal or differ for both cases, we note that the scaled variance compared to the scaled mean squared 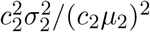is also the same. However, both cases differ in their values of 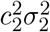 which seem to play a negligible role. The mRNA excursions and their strength directly translate to the protein level (lower panels of 4E). We have thus explained the behaviour of the protein mean for the pair of high variance and medium correlation. For the other pair with high correlation and medium variance a similar behaviour is not observed, compare figure 4F and G. The lack of this behaviour hints at the crucial role of a sufficiently large environment variance in order to expose relevant excursions.

**Figure 7.**
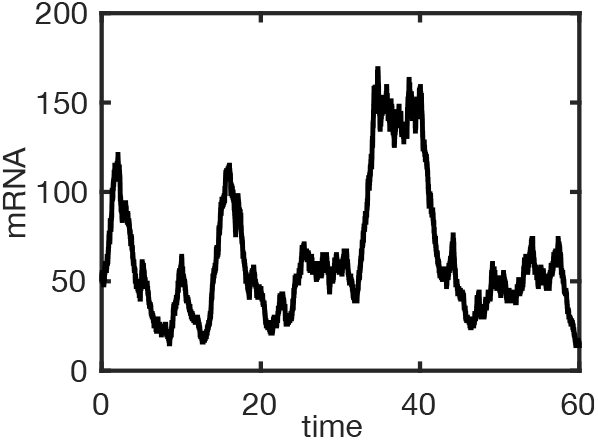
A sample trajectory of the mRNA counts for the approximate marginal simulation, i.e., the analog to fig 2H

**Figure 8.**
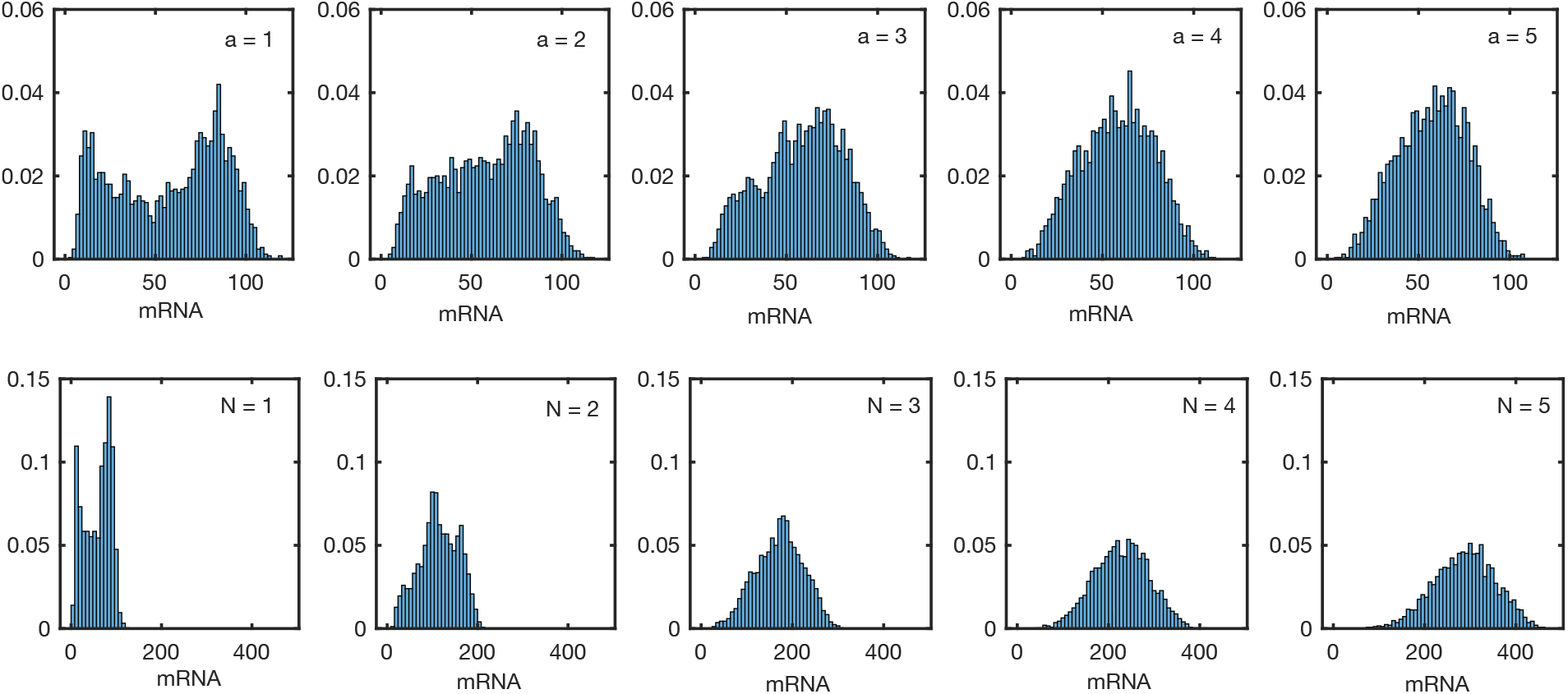
Details on promoter-mediated transcription. Histograms of the stationary mRNA, simulated from the exact system using the Doob-Gillespie algorithm. Final time point was t = 60. For the upper row, the transition rates were multiplied by a = 1, …, 5. For the lower row N was increased from 1 to 5. Otherwise, the parameters were as in 2E. The number of trajectories was 2500.

**Figure 9.**
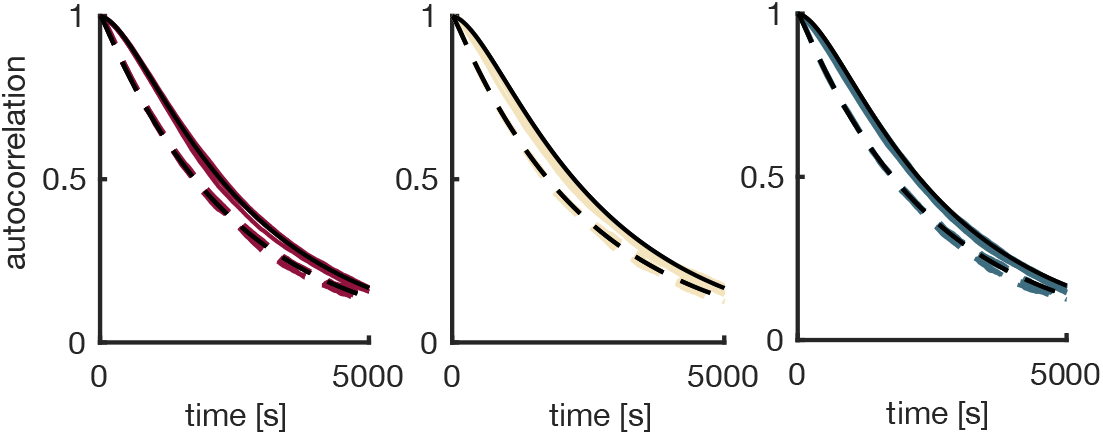
Details on the gene expression model in the fast environment. The protein autocorrelation as a function of the lag (dashed = fast mRNA timescale, solid = slow mRNA timescale). Analytical autocorrelation for the reference model (black), shown with the correlation coefficient of protein trajectories for the embedded model (coloured) at different conditions (from left to right: varied modulation by Z_2_, Z_3_, Z_4_), estimated from 10000 trajectories. It was computed from the last time point, paired with previous time points.

**Figure 10.**
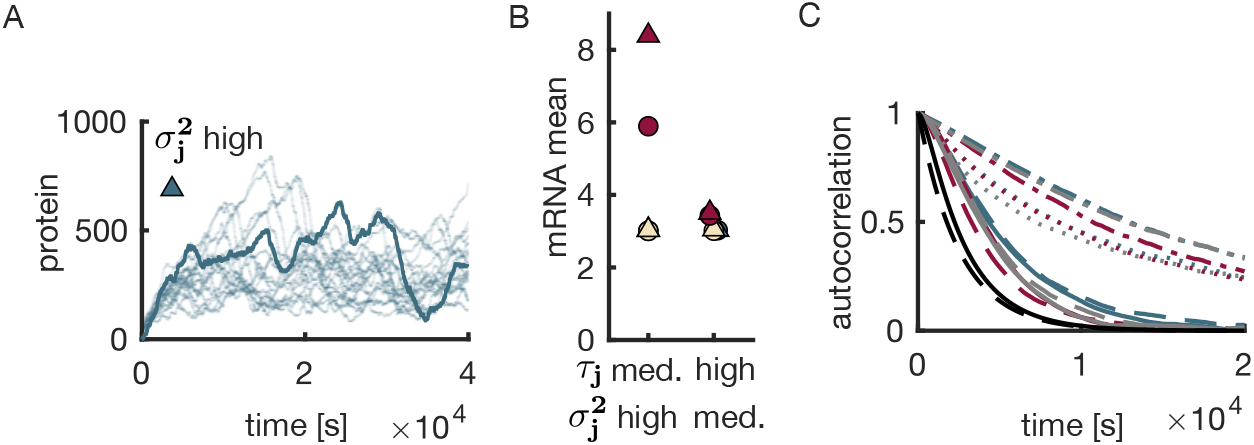
Details on the gene expression model in the slow environment. A. Sample trajectories of the protein for the maximal σ^2^ value and medium (protein-like) correlation time τ_2_ with the fast mRNA timescale. B. The mRNA mean at time t = 40000 estimated from 10000 sampled trajectories for the different conditions. Different colours for the same tick were shifted horizontally to improve readability. C. The protein autocorrelation as in 4D for all conditions in the medium and high regime.

When we now turn towards the variance (fig 4C), we see a very different behaviour compared to the regime of low correlation (fig 3D). The variance for the low regime was dominated by the base variance and a contribution by the product of *τ*_*j*_ and 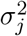, which increased as *τ*_*j*_ approached the mRNA timescale and 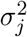 its maximal value. For the medium and high regime, we see no significant contribution of the base variance any more. It is still noticeable when we look at the differences between the slow and the fast mRNA time scales for an R_3_ and an R_4_ modulation. However, what seems to become the dominating factors are the mean level of the protein (largely caused by excursions) and the environment variance. Consequently, the trajectories, figure 4E, look qualitatively different from the reference case.

While for the protein mean and variance, the environment variance 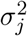played the essential role, this is not the case when we look at the autocorrelation of protein trajectories, figure 4D. We analytically computed the autocorrelation function for the reference system as in [54], and compared it to the correlation coefficient obtained from 10000 Monte Carlo sample trajectories. We computed the correlation coefficient between the final time point at *t* = 40000 and time points further in the past, increasing the lag from 0 to 20000. To avoid cluttering, we excluded the R_3_ modulation. Its behaviour is qualitatively the same as R_2_ and R_4_, see appendix, figure 10C. Both the medium and the high regime differed significantly from the reference case, showing a slower autocorrelation decay. When comparing the high regime and the medium regime, the high regime even showed a much slower decay than the medium regime. When we look for differences among the modulation by R_2_ and R_4_ combined with the slow and fast mRNA timescale, we find that the R_4_ modulation in the fast mRNA timescale shows the slowest decay, and the R_2_ in the slow regime the least slow one. When looking at sample trajectories, figure 4F and G, we observe that two individual trajectories can remain separate for a rather long time. For the R_4_ modulation in the fast mRNA timescale, we see that the trajectories show very little mixing behaviour (fig 4G). This illustrates the strong correlation between even far apart time points.

To summarize, we observed an increase in the protein mean, when both the correlation time and the variance of an environment component that modulates a decay reaction, are large enough. We attributed this effect to excursions. We saw a stronger effect for the modulation of the mRNA decay than the protein decay. Interestingly, our findings differ from Keizer et al. who investigated the same gene expression model in a different environment, i.e., using log-normally distributed slow extrinsic noise. For the parameter regime considered in their work, the mean protein levels were the same regardless of modulating the mRNA decay or the protein decay. In our case study, the increase of the mean contributed to an increase in the variance, which, for the regime of medium to high autocorrelation, is no longer dominated by the base variance and 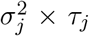. The protein autocorrelation increases essentially with the autocorrelation of any environment component, with minor differences for the considered conditions.

### 3.3 First- and zeroth-order modulation by a correlated environment

While in the two previous case studies we investigated a multi-component correlated environment targeting the same reaction and uncorrelated environment components targeting multiple reactions, we now look at correlated environment components targeting multiple reactions. We choose a system for which we computed the stationary mean in our recent work [51]. Hence, we can compare the results of the approximate marginal simulation to the exact one. Let us start by describing the system, see figure 5A. A birth-death process *X* is embedded into an environment which modulates its death reaction. The environment component *Z* is itself a birth-death process. In phases of *Z* = 0, molecules of *X* accumulate, which we call excursion. When the environment switches fast, these periods are negligible, but if it is slow, these excursions, although rare, can dominate the stationary mean. In the previous case study, we have observed that the tilted Hawkes model can generate excursions. With analytic expressions for the stationary mean at hand in the present case study, we aim to test whether the impact on the stationary mean is also captured quantitatively.

To mitigate the effect of excursions, a controller component *U* is added. It senses the environment value *Z* and takes action by down-modulating the birth rate of *X*. The sensing is realized by birth reactions of *U* that are modulated by *Z*. Independently of *Z*, the *U* molecules degrade. Then it depends on whether the controller reacts fast enough to flatten excursions. Formally, the joint process (*U, Z*) is used as the environment for the birth-death process *X*, hence this is an example of a birth-death process embedded in a correlated environment. The control goal is to stay close to the set point, which corresponds to the homogeneous system with averaged environment and controller. For two different controller speeds, the figure 5B depicts the stationary mean as a function of the environment speed. When the environment is fast, independent of the controller, the set point goal is achieved. If the environment gets slower, it is not surprising that the faster controller is more successful with attaining the control goal. As a function of the environment speed, the stationary mean has the maximal point in the interior, which we explained in [51]. The tilted Hawkes model captures this behaviour. In the sample trajectories figure 5C and D of the state estimates, we notice that the fast controller tracks the environment and the slow one only follows with a sincere delay or not at all. Hence, the state estimate mimics capture the behaviour of environment and controller qualitatively. We note here that we did not encounter negative values of the state estimates 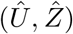.

Returning to the slow controller in figure 5B, we observe a deviation of more than 100% from the set point, which the tilted Hawkes model does not exhibit. To see that the excursions cause the deviation from the set point to a large extent, we examined the shares that the different environment states contribute to the stationary mean (fig 7d in [51]). Also, when we compare sample trajectories of the exact system (fig 5E) and the tilted Hawkes model (fig 5F), we see an excursion occurring for the exact system at around *t* = 150, while the tilted Hawkes model does not show an excursion. We conclude that the tilted Hawkes model underestimates the effect of the excursions on the stationary mean. We did not rule out that for a slower environment, the severe deviation from the set point and the rise to a plateau value for the slow controller can be qualitatively captured. Also, it is not unlikely that the simulation time was too short for the last confidence interval (slowest considered environment, slow controller) in figure 5E to properly capture the mean. When we cross-checked the Monte Carlo estimation procedure by the exact system with the same hyperparameters, the mean was often not captured by the confidence interval. This supports the hypothesis of failed convergence. Beside this possible explanation for the discrepancy between the exact mean and the approximation, we recall that in the previous case study, we saw the excursions only for sufficiently high values of *σ*^2^. Consulting these findings, we hypothesize that the stationary variance of *Z* could be too small for excursions to arise. In summary, we see that, on the one hand, the tilted Hawkes model captures important features of the model such as the controller (not) following the environment and the maximal deviation occurring in the interior environment speed. The quantitative deviation attributed to underestimating excursions, on the other hand, poses a clear limitation to the approximation.

## 4 Conclusion

In this paper, we recycled an approach of optimal linear state estimation for the doubly stochastic Poisson process for the use in approximate marginal simulation of CRNs in random environment. The Hawkes approximations studied here provide a convenient way to add variance to the system, while using interpretable parameters, i.e., the mean, covariance and auto/cross-covariance decay of the environment. This matches well with the situation of the experimentalist who has limited information about the environment, but would like to include that information in the model. Reversely, the partial observations of the system that is being modulated could be used to infer the three environment characteristics using the present Hawkes approximation. For this purpose, it would be desirable to derive path likelihoods for the Hawkes modelled CRNs. At the same time, it is a limitation of our approach that the environment is restricted to the exponentially decaying auto/cross-covariance. Being especially well-tailored for linear environment that modulate zeroth-order reactions, we expect that our Hawkes model can be used to replace linear leafs in the reaction network graph and linear transit networks. As a word of caution, we would like to recall that for this kind of model reduction to work the replaced linear environment must not be too discrete or bimodal, as we have seen in the first case study.

We emphasize that the Kalman-like approach does not require a diffusion approximation in order to be applied. With the link being a shared integral equation for the estimation kernel, the results from the continuous-time additive white Gaussian noise channel could be transferred. While its original purpose was state estimation, we recycle the approach here to use it for approximate marginal simulation. A related linearizing approach, that is sometimes taken, models the environment components as Ornstein-Uhlenbeck processes, or equivalently chemical Langevin equations. For this approach, one can criticise that the intensity can become negative. For our approach, when using independent environment components, positivity is guaranteed by the one-dimensional Hawkes model. For a correlated multi-component environment that modulates a single reaction, we derived a sufficient criterion for positivity. However, in general, we could not guarantee that the filter remains positive, which restricts the approach. We anticipate, however, that the possibility of negative rates is less restrictive than for the Ornstein-Uhlenbeck process, because we use marginal rates. The latter ones have a smaller variance, and thus explore a smaller range of intensity values, making negative intensities less likely. To rule out the possibility that the filter becomes negative, a different projection criterion could be applied. Atar and Weissman showed that the natural loss criterion provides a principled choice that is well-suited for positive random quantities [3]. In figure 6 we illustrate how the optimal Hawkes parameters are altered when the natural loss criterion is used.

The remark after lemma 2.2 hints at a future application for model reduction. A matched covariance measure can also guide the design of approximations for non-linear systems. It is the second term when expanding the logarithm of the characteristic functional of the random measure associated with the counting process in terms of cumulants [14, §9.5, proposition 9.5.V]. Higher order terms can successively be matched.

Our characterization theorem 2.5 sheds light on the moment closure 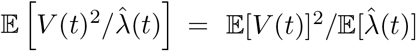, which was misconceived before. In [57] this moment relation and its supposed equivalence with 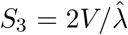 was erroneously used to derive the Gamma filter from the CIR-modulated reaction channel. This would have justified the Gamma filter as being exact for the CIR-modulated counting model. However, we showed that the moment closure instead imposes the optimal linear filter. In [50], we compared the Gamma and optimal linear filter for the estimation of the mutual information in the two-state model of gene expression. We noted that the optimal linear filter improves the computation time with no decrease in accuracy.

The applications for which either of the two approximate filters is preferable need still to be determined. The characterization result can be used to guide the intuition.

In the second case study, we investigated a two-stage gene expression model in a random environment. Each reaction was modulated by a separate environment component stochastically independent of the others. In the simulation study, we varied the second-order statistical properties of only one environment component at a time, while leaving the other components at a negligible level. We saw the most noticeable effect for decay reactions modulated by slow environments with a large variance. The stationary protein mean was largely increased. For the parameters studied, the effect was much stronger for the mRNA compared to the protein decay modulation. The mRNA stage was both upstream and faster. Two possible explanations can be considered. An environmental effect in the upstream component dominates over one in the downstream component, or an environmental effect in the faster stage dominates over the slower stage. We argue that the stronger effect for the mRNA modulation is not a consequence of being in the upstream layer, but a consequence of the faster mRNA timescale. This would in line with the different effect for different mRNA timescales that we observed and explained by investigating the mRNA level. We propose to check this claim by inverting the timescales of the upstream and downstream layer. For the variance, we identified various factors that increase it. They dominate for different environment correlation regimes. In particular, for the regime of low correlation, the dominating factor on top of the base homogeneous variance was the product of variance and autocorrelation. This is precisely the term that complements the Poisson variance in the Markov modulated Poisson process, i.e., the simplest zeroth-modulation case. We speculate that the variance can be approximately decomposed in an analogue of proposition 2.4 for first-order modulation. For the transfer of variance to downstream components via synthesis reactions, we speculate that results like proposition 2.4 can give a hint, even if the synthesis reaction is modulated in addition.

A natural next question to ask is what happens if two environment components are simultaneously changed. If both *Z*_2_ and *Z*_4_ have high autocorrelation and high variance, will we observe a Synergistic behaviour of the excursions? Do the effects add up, or do they multiply? This is a subject of future research. Finally, we observed that the Hawkes approximation was able to capture the effect of mean deviation due to excursions. However, when comparing it to the exact numeric values in the third case study, we saw that it underestimates the effect. It would be worthwhile to investigate in what way it overestimates how large the parameters must be for the effect to become apparent.

In some cases, the Hawkes modelling leaves room for alternative partitions into subnetwork and environment. In the third case study, we investigated the stationary mean for a birth-death process *X* in an environment consisting of a death rate modulating environment component *Z* and a controller *U*. While we applied the tilted Hawkes model to the environment (*U, Z*), we could alternatively define *Z* alone as the environment and (*U, X*) as being modulated by it.

Although several approximate filter schemes have been proposed in the last decade, there has been limited progress on a systematic study of their structural properties and estimations of their accuracy. For the optimal linear filter, we provided structural results in terms of the second-order moment agreement for zeroth-order modulation. We anticipate that the optimal linear filter can be used as a base case which can be adjusted and improved. We advocate that adjustments should be done in a controlled and hierarchical way to account for structural properties at each level of adjustment. We imagine that a principled approach for adjustments can take at least two routes. First, a canonical adjustment is the incorporation of increasing orders of dependency on the trajectory of reaction counts. We covered the case of first-order dependence, the next step could be a second-order dependence. A second route could use variational methods to find the optimal functional form of improvements, similar to [43]. Thereby, we would not impose the polynomial order, but control for the trajectory features that are taken into account. For instance, this could be hierarchically achieved by an increasing family of sigma-algebras.

## Appendix A. Detailed calculation of the covariance matrix

We get

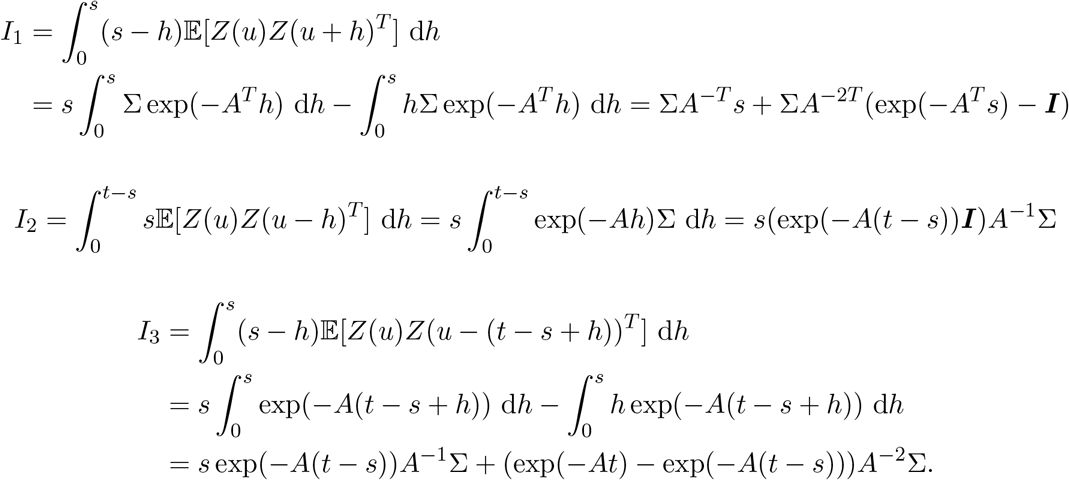

## Appendix B Details on the case studies

### B.1 Invertibility of matrices for the promoter-mediated transcription

This section discusses how to make *A* and 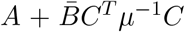 invertible. The matrix *A* in Eq. (3.1) has vanishing column sums, i.e., **1**^*T*^ *A* = **0**^*T*^ for the vector **1** = [1, …, 1]^*T*^. For this reason *A* is non-invertible. The matrix σ^2^ has column and row sums zero, i.e., **1**^*T*^ σ^2^ = **0**^*T*^ and σ^2^**1** = **0**. We derive an equation equivalent to Eq. (1.7) -Eq. (1.8) for the reduced state 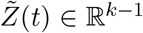, droping the *k*-th coordinate of *Z*(*t*). This way, we make *A* invertible by reducing its dimension.

To this end, we define Γ and decompose the matrices A and σ^2^ as

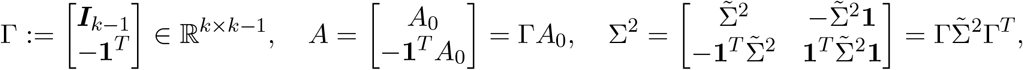

with 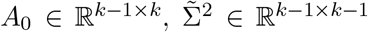. Then we define 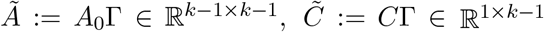. Let 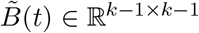 solve

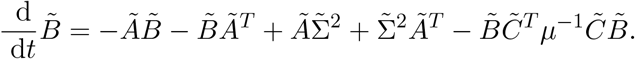

Then we derive the evolution equation for

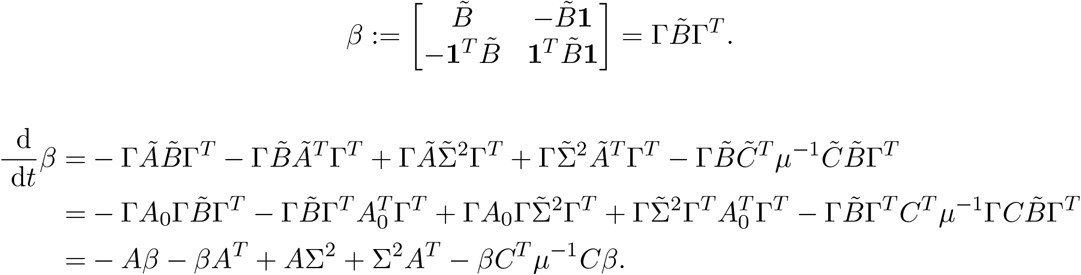

Hence *β*(*t*) = *B*(*t*) for all *t ≥* 0. Let further 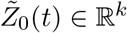 satisfy

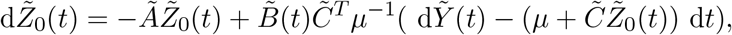

where the intensity of 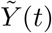 is assumed to be 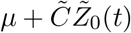. Then 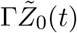 satisfies

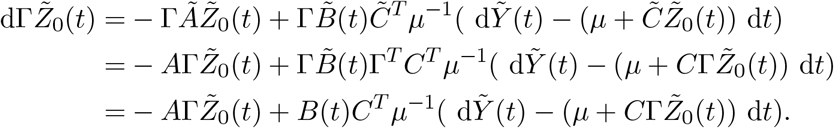

Hence for 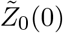equal to the first *k* − 1 entries of *Z*_0_(0), we have 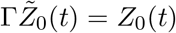 for all *t ≥* 0 and in particular 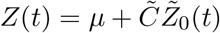,hence 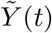 is equal to *Y* (*t*) in distribution.

## Acknowledgments

MS would like to thank Maleen Hanst and Felix Reinhardt for their consent to use their code structure as part of simulating the case study 1, Nikita Kruk for providing the code to simulate the case study 3 and Christian Wildner for general discussion.

